# Chronic stress promotes sex-divergent Type 1, 2 and 17 responses in brain resident lymphocytes

**DOI:** 10.64898/2026.08.31.748329

**Authors:** Simon Paris, Ke Cao, Yoobhin Park, Kevin Junwon Lee, Sicong Zhou, Deeva Uthayakumar, Miguel Novelo, Alara Tuncer, Chenxi Qian, Chao Wang

## Abstract

T helper cytokines such as IFNγ and IL-17 are dysregulated in a subset of patients with depression and are associated with poor responses to treatment. Both IFNγ and IL-17 have been shown to regulate neural circuits involved in behavioral changes implicated in depression. However, the cellular source and the regulatory mechanisms governing these cytokines in this context remain incompletely understood. We analyzed brain lymphocytes in response to unpredictable chronic mild stress (UCMS), a model of depression with strong face and construct validity. We confirmed that UCMS induced depressive-like behavior in both male and female mice. Interestingly, UCMS preferentially reduced regional synaptic density in female and neuron counts in male hippocampi. Using single cell RNAseq, we identified T helper cytokine signatures to be enriched in distinct lymphocyte lineages expressing tissue resident markers and are regulated by both sex and chronic stress. Female brain-resident γδT cells are significantly enriched for type 2 and type 17 signatures and have higher expression of inflammasome genes such as Nlrp3 and Il1b whereas male αβT cells and NKT cells are enriched for type 1 signatures and have elevated Cxcr3 and Il27ra expression. Chronic stress amplified the sex differences and preferentially increased the type 1 response including IFNγ production in males at both the transcriptome and protein level. Type 17 response (e.g. IL-17A and IL-17F production) is also increased in response to UCMS, however with sex-divergent polarization: IL-22 is decreased in female and increased in male brain lymphocytes. Mechanistically, we found that UCMS suppressed P4ha1 in male and Zfp36 in female brain tissue resident lymphocytes. P4HA1 was previously shown to restrict mitochondrial function and limit type 1 responses in CD8 T cells, whereas ZFP36 is a suppressor of inflammatory cytokine translation. Consistent with this finding, ex vivo CD3-dependent stimulation of brain immune cells showed stress-dependent increase in cytokine production with sex-specific effects. Our study provides the cellular and molecular framework for chronic stress-induced tissue resident lymphocyte dysregulation and its sex bias in the brain, with implications for depressive disorders marked by inflammation.

## 1. Introduction

Immune cell dysfunction is associated with chronic stress, notable behavioral changes and increased risks of developing major depressive disorder. Epidemiological evidence supports that at least a subset of depression patients present with an increase in broad T helper cytokine profile including both proinflammatory and anti-inflammatory cytokines (e.g. IL-17, IL-10, IL-9, IFNγ, IL-5, IL-13, IL-2) underlying general immune dysregulation (Syed et al. 2018). Interestingly, an increase in IFNγ and IL-17, but not type 2 cytokines, is associated with non-responsiveness to anti-depressant treatment (Syed et al. 2018), alluding to their possible role in disease pathogenesis. Indeed, previous work demonstrated functional roles for IFNγ and IL-17 in regulating neural circuits in steady-state mice influencing social interaction and anxiety, both of which are commonly dysregulated by depression (Filiano et al. 2016; Alves de Lima et al. 2020). Although an increase in IL-17- and IFNγ-producing T helper 17 cells was reported in the peripheral blood of some MDD patients (Schiweck et al. 2020; Ghosh et al. 2020), the cellular sources of these cytokines within the brain remain poorly defined in both depression and relevant stress models.

T helper cytokines such as IL-17 and IFNγ can derive from various immune cell lineages such as conventional T cells, Innate Lymphoid Cells, gamma-delta T cells (γδT), Natural Killer (NK) cells, NKT cells and mucosal-associated invariant T (MAIT) cells (Fan and Rudensky 2016). Previous work using the learnt helplessness model, which induces physical stress, identified CD4^+^IL-17^+^ cells in the brain (Beurel et al. 2013). In this model, brain resident CD4 T cells are dependent on the presence of the gut microbiome (Medina-Rodriguez et al. 2023). In the social defeat model, which induces psychological stress, IL-17-producing γδT cells were found to expand in the intestine and accumulate in the brain leading to exacerbated depressive-like behaviors (Zhu et al. 2023). As both studies used male mice and given the pronounced sex bias in depression and stress physiology, it is unclear whether similar immune changes occur in females under stress. Furthermore, not all stressors elicit identical physiological outcomes. Distinct immune responses can arise depending on the neural circuits engaged by the type (physical or psychological), duration (acute or chronic) and predictability of stress (Haykin and Rolls 2021).

The unpredictable chronic mild stress (UCMS) model is a widely used model of social and environmental stress paradigm that induces depression-associated brain pathology, behavioral deficits, and is routinely used to test antidepressant efficacy (Mineur et al. 2006; Willner 2017; Planchez et al. 2019; Rawat et al. 2022). Hippocampal neurons are targeted by stress (McEwen 1999) and are obligatory targets of fluoxetine and ketamine, two commonly prescribed selectivw serotonin reuptake inhibitor and N-Methyl-D-aspartate receptor antagonist respectively, for their anti-depressive effects (Shuto et al. 2020; Rawat et al. 2022). Previous work demonstrated that UCMS induced functional impairment of neuron synapses in the hippocampal CA1 region (Kallarackal et al. 2013) and decreased neurogenesis (Du Preez et al. 2021). UCMS was also shown to induce significant changes in other brain regions relevant to depression such as prefrontal cortex (Musaelyan et al. 2020) and amygdala (Vyas et al. 2002). Although the UCMS model demonstrates robust behavioral and neurobiological activity, a comprehensive characterization of its immune components is still lacking. Chronic variable stress (similar to UCMS but with shorter duration) activates hematopoietic stem cells (Heidt et al. 2014), epigenetically reprograms monocytes towards an inflammatory phenotype (Barrett et al. 2021) and impairs iNKT cell function (Rudak et al. 2021). In the brain of UCMS mice, male microglial (Iba1^+^) cells were found to increase in various brain regions (e.g. cortex, amygdala, hippocampus) (Troubat et al. 2021). Here we characterized both female and male C57BL/6 mice subjected to UCMS, focusing on stress-induced changes in brain lymphocyte responses using flow cytometry, single cell RNAseq and cytokine profiling.

## 2. Material and Method

### 2.1. Mice

Male and female C57BL/6 mice were bred in house and applied to experiments between 8-24 weeks of age. All mice are grouped by sex and co-housed at the time of weaning. Age- and sex-matched littermates were used for each comparison group in all experiments. All mouse experiments followed guidelines outlined and animal user protocols approved by the Animal Care Committee at Sunnybrook Research Institute in Toronto, Ontario, Canada.

### 2.1. Unpredictable chronic mild stress (UCMS) model

Littermate mice were either left unstressed or exposed to a shuffled schedule of daily mild stressors for up to 8 weeks. Stressors are scheduled in random order each week to reduce habituation, a glucocorticoid-dependent process to self-regulate long-term deleterious effects of stress (Herman 2013). Stressors include: empty cage (4 hours), tilted cage (3 hours), shaking (2 hours), wet cage (4 hours), fasting (food only, 24 hours), restraint (2 hours) and dark/light cycle disruption (48 hours) following a published protocol (Frisbee et al. 2015; Burstein and Doron 2018). Mice were returned to home cage after each stressor. To monitor progress of chronic stress, mice were weighed weekly. Behavioral tests were performed once per mouse typically at the end of the UCMS paradigm. See **Table 1** for full UCMS protocol schedule description with stressors and behavioral test. A total of 140 animals were used for the presented study. Serum corticosterone levels were measured, typically at the end of the UCMS paradigm, using a competitive ELISA Kit (Thermofisher, Cat. EIACORT). Euthanasia: at the end of the UCMS protocol, mice were placed in an incubation chamber and exposed to gradual and lethal levels of CO_2_. At the cessation of breathing, cervical dislocation was performed. The chest cavity was exposed and a 27G needle was inserted into the left ventricle for intracardial perfusion with and PBS and optional second perfusion with 4% paraformaldehyde for brain fixation when needed.

**Table 1.**
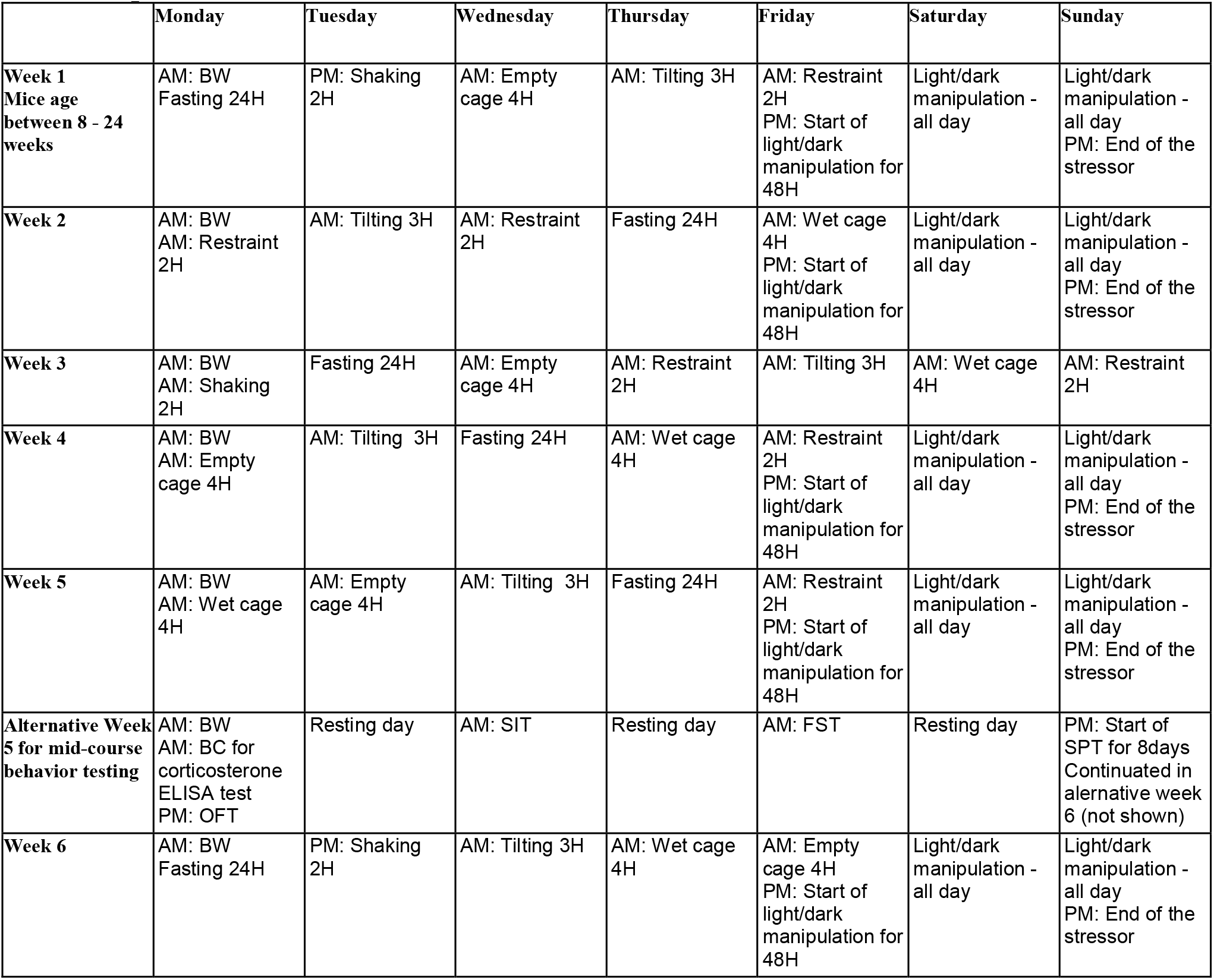
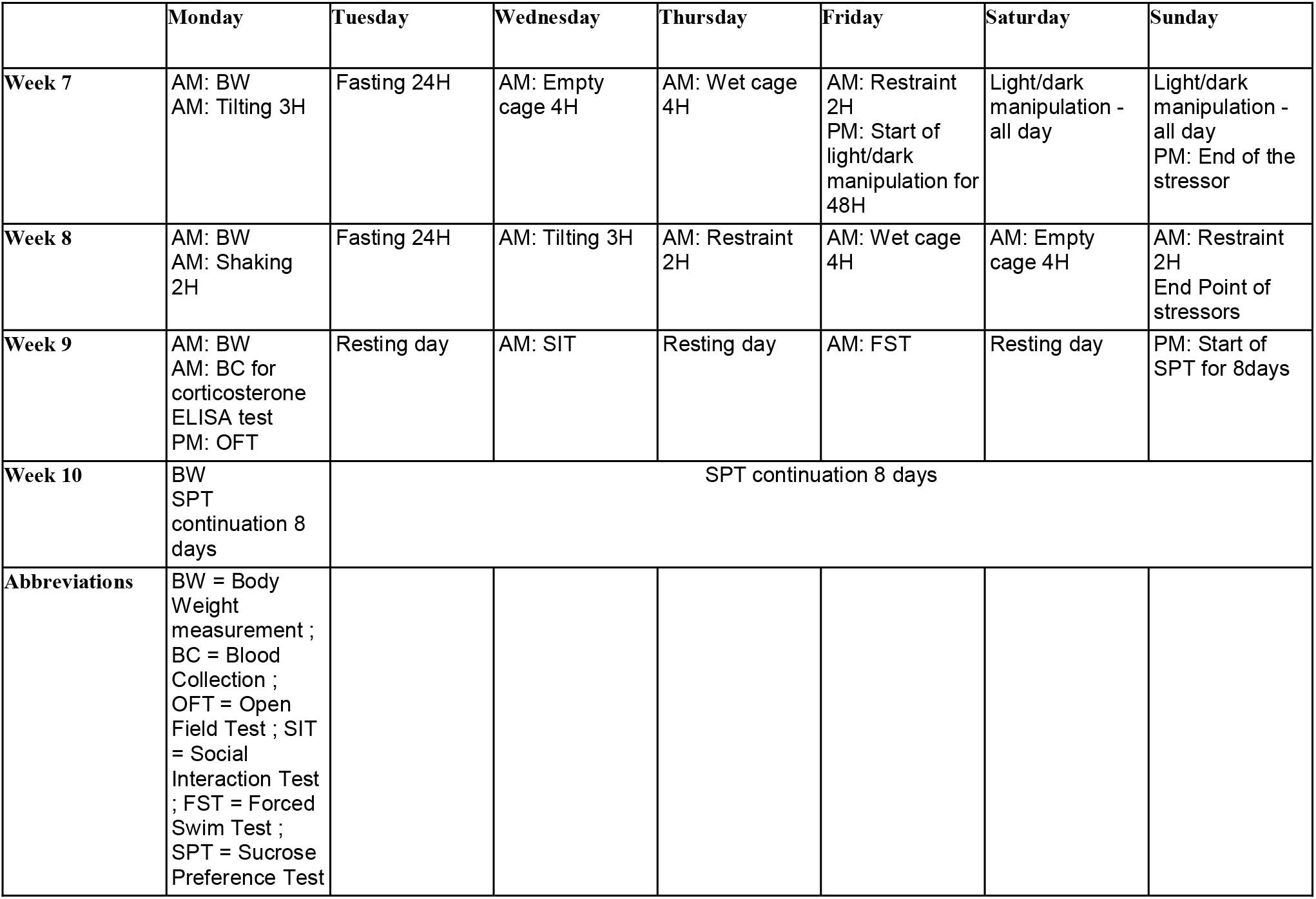
Unpredictable Chronic Mild Stress model schedule and behavioral tests.

### 2.2. Behavioral tests

Behavioral tests were performed in specialized behavioral rooms at the Sunnybrook Vivarium with proper acclimatation to the procedure room for a minimum of 60 minutes prior to starting the tests. The Open Field Test (OFT) was used to assess anxiety similarly to (Seibenhener and Wooten 2015). Mice were placed in the SuperFlex Open Field system (Omnitech Electronics, Inc.) for 10 minutes and movement was captured by infrared beams. Data were analyzed by Fusion software. The level of anxiety is calculated by the time spent in the outer zone of the arena. Social interaction test (SIT) was performed as described in (Lopatina et al. 2014) in the same SuperFlex open field system. Briefly the social interaction preference of the test mouse was determined by the time it spent with another mouse placed in a wired cup in the open field and data analyzed by Fusion software. Forced Swim Test (FST) was performed similarly to (Can et al. 2012). Briefly, mice were placed in lukewarm water (24°C) and allowed to move freely for 6 minutes and movement captured by an overhead camera. Ethovision software (Noldus) was used to analyze immobility in the last 4 minutes of each session. Sucrose preference test (SPT) was performed as described in (Liu et al. 2018) in a 8 day protocol. SPT measurement is calculated as % of sucrose consumption divided by the sum of total liquid consumption (water + sucrose).

### 2.3. Western blot

The expression levels of PSD95, Synaptophysin (SYN) and Myelin-basic protein (MBP) in the mouse hippocampus were determined by western blot analysis.

#### Tissue preparation and protein extraction

Following UCMS protocol, mice were euthanized and transcardially perfused with PBS. Flash frozen hippocampi were resuspended in RIPA buffer with protease and phosphatase inhibitors for total protein extraction. Tissue dissociation was achieved manually by pipetting and sonication. Samples were clarified by centrifugation at 16,000xg at 4°C for 20 minutes. Protein quantification was assessed using the Pierce™ BCA Protein Assay Kit (Thermo Fisher Scientific, cat. 23225).

#### SDS-PAGE

Sample volumes are normalized based on their protein concentration and proteins were resolved on Bolt 4-12% Bis-Tris Plus mini protein gels (Thermo, cat. NW04120BOX). After SDS-PAGE gel electrophoresis samples were transferred on low fluorescence PVDF membrane 0.2μm pore. After transfer, membranes were stained for total protein quantification using the Revert™ 700 Total Protein Stain Normalization Protocol (Licorbio, cat. 926-11015). Total protein signal on membranes was analyzed on a Licor Odyssey F instrument using the 700nM channel (**Supplementary Figure 2**). Membranes were then washed to go on with the regular antibody staining. After blocking with 5% BSA in TBS-Tween 20 (TBST) buffer for 1 hour at room temperature and washed 3 times for 5 minutes in TBST buffer, the membranes were incubated overnight at 4°C with primary antibodies (anti-PSD95 from Synaptic Systems Antibodies, cat. 124 008, dilution 1:500; anti-Synaptophysin from Invitrogen, cat. MA5-14532, dilution 1:200 or anti-MBP from Sigma-Aldrich, cat. MAB382, dilution 1:200). The following day, membranes were washed 3 times with TBST buffer and then incubated with secondary antibodies in the dark for 1 to 2 hours at room temperature (Goat anti-rabbit AF488 from Invitrogen, cat. A11008, dilution 1:500 or Donkey anti-mouse AF647 from Invitrogen, cat. A-31571, dilution 1:1000). Fluorescence signals were analyzed on a Vilber Fusion Fx imager. Image J was used to quantify the integrated densities of protein-specific ROIs (LUT inversion and subtraction of background) of each band before being normalized using the total protein quantification method. In quantitative Western blotting (QWB), normalization mathematically corrects for unavoidable sample-to-sample and lane-to-lane variation by comparing the target protein to an internal loading control. The internal loading control is used as an indicator of sample protein loading, to correct for loading variation and confirm that observed changes represent actual differences between samples.

### 2.4. Immunofluorescence and microscopy

#### Brain slices preparation and immunofluorescence

Mice were transcardially perfused with PBS, followed by 4% paraformaldehyde (PFA). Brains were isolated, post-fixed for 24□h in 4% PFA and cryoprotected in 30% sucrose solution until full tissue submersion (∼ 48 h to 1 week). Tissues were embedded in the O.C.T compound (PANTek Technologies LLC, cat. 23-730-571) and stored in −80℃ until ready to use. 30µm-thick coronal brain slices were sectioned using the Leica CM3050 S cryostat and stored in a 96-well plate with 200µl PBS. Immunostaining was performed in 24-well plates, where free-floating sections were blocked using 10% donkey serum (Jackson ImmunoResearch, cat. 017-000-121) and 0.1% triton-X 100 (Sigma-Aldrich, cat. X100-500ML) for 1 h. Brain sections were incubated overnight at 4℃ with 5% donkey serum, 0.1% triton-X 100, anti-mouse NeuN (Sigma-Aldrich, cat. ABN90, dilution 1:500). Unbound primary antibody was washed 3 x 5 mins with PBS. Secondary antibody, donkey anti-guinea pig AF594 (Jackson ImmunoResearch, cat. 706-585-148, dilution 1:1000) was incubated in the dark with 5% donkey serum and 0.1% triton-X 100 at room temperature for 1 h. After 2 x 5 mins of wash with PBS, sections were mounted on VistaVision™ HistoBond® Premium Adhesion Slides (VWR, cat. 16004-406) using ProLong™ Diamond Antifade Mountant with DAPI (Invitrogen, cat. P36971) and Fisherbrand™ Cover Glasses (Thermo Fisher Scientific, cat12-544-GP).

### 2.5. Confocal Microscopy

#### Image acquisition

Digital images were scanned using a Nikon A1 confocal microscope (Nikon Instruments Inc., Tokyo, Japan) equipped with a 20× objective lens. Z-stack images were acquired at 1.625 μm intervals to capture the full thickness of the immunostained sections. For all regions, quantification was performed within a 0.62 mm × 0.62 mm field of view.

#### Image processing for quantification

Quantitative analyses were conducted on male and female mice from the unstressed and 8-week UCMS experimental groups (n = 3 mice per group). Regions of interest (ROIs) for this study included the following hippocampus regions: CA1, CA2, CA3, subgranular zone (SGZ) of dentate gyrus (DG), suprapyramidal blade of DG (sDG) and infrapyramidal blade of DG (iDG). Images were collected from coronal brain sections corresponding to Bregma −1.555 mm to −1.755 mm, based on the Allen mouse brain atlas. Imaging parameters, including laser intensity, gain, and offset, were maintained constant across all samples within the same sex and ROI to allow accurate comparison of fluorescence intensity. Parameters were adjusted as needed between different ROIs and between sexes to account for region- and sex-specific signal variability. Neuronal layer thickness in the hippocampal CA1, CA2, CA3, sDG and iDG regions as well as the neuron count within the SGZ were evaluated using the z-stack maximum intensity projection images. Nikon NIS-Elements AR software (Nikon Instruments Inc., Tokyo, Japan) was used. The height of the cell layer at three fixed, equidistant locations within each field of view were used to measure the pyramidal layer thickness. For the number of neurons within the SGZ, manual counting was performed by counting DAPI+NeuN+ cells within the given field of view. Neuron count in the CA1, CA2, CA3, sDG and iDG regions of the hippocampus was quantified using QuPath (QuPath, v0.6.0). Analyses were conducted from three representative optical planes of 1.625μm-thick z-stack capturing the whole ROI were selected for analysis in each section. Three sections that are approximately 60μm apart were assessed and the average used for calculations. Within each photomicrograph, neuronal counts were obtained from three equidistant regions using a 100 μm × 100 μm sampling grid for the CA1, CA2, and CA3 pyramidal layers, and a 50 μm × 50 μm sDG and iDG. Manual cell counting of DAPI+NeuN+ cells were performed. Statistical analyses for neuronal layer thickness and neuron count were conducted via GraphPad Prism version 10.0.0 for Mac (GraphPad Software, Boston, Massachusetts USA).

### 2.6. Stimulated Raman scattering (SRS) microscopy

A picosecond laser system (picoEmerald, Applied Physics & Electronics) served as the light source for SRS microscopy. It generated a 2-ps pump beam (tunable between 770–990 nm, bandwidth 0.5 nm, corresponding to ∼7 cm⁻¹) and a Stokes beam (1031.2 nm, bandwidth ∼10 cm⁻¹) at 80 MHz. The Stokes beam was modulated at 20 MHz using an internal electro-optic modulator. Both beams were spatially and temporally aligned and directed into an inverted multiphoton laser scanning microscope (FV3000, Olympus), where they were then focused onto the sample using a 25× water-immersion objective (XLPLN25XWMP, 1.05 N.A., Olympus). The transmitted pump and Stokes beams were collected with a high-N.A. oil-immersion condenser (1.4 N.A., Olympus) and passed through a bandpass filter (893/209 nm BrightLine, 25 mm, Semrock) to block the Stokes beam. The pump beam which passed through the filter was detected by a Silicon photodiode (10 × 10 mm, S3590-09, Hamamatsu) biased at 64 V DC to raise the saturation limit and shorten the response time. The photodiode output was terminated with a 50 Ω load and pre-filtered with a 19.2–23.6 MHz bandpass filter (BBP-21.4+, Mini-Circuits) to minimize the laser and scanning noise. The resulting signal was demodulated by a lock-in amplifier (HF2LI, Zurich Instrument) at the 20 MHz modulation frequency. The in-phase output was sent to the microscope’s analog input (FV30-ANALOG, Olympus).

#### Myelin imaging

For imaging the myelin, characterized by the CH2 vibrational mode at 2940 cm⁻¹, the pump laser was tuned to 797 nm. The measured laser powers on the sample were 40 mW (pump) and 100 mW (modulated Stokes). 12-bit images were acquired using Olympus Fluoview 3000 software, and volumetric datasets were obtained by z-stack imaging with 1 µm step size along the z-axis. Image acquisition was set at 4μs pixel dwell time and 2048*2048 frame for CA1 region and 1024*1024 for CA3 and Dentate Gyrus, which costs a total of 17 seconds per frame and 151 seconds per stack. Myelin was found to accumulate in the molecular layer and hilus region instead of the granule cell layers. Therefore, we targeted our quantification region at the nearest region next to the CA1, CA3 and Dentate Gyrus. Flat-field correction was done using the BaSiC Plugin in Imagej. Max projection images were created from the stack images using Imagej. Myelin was segmented by thresholding and the average intensity in each hippocampal region was calculated.

### 2.7. Flow cytometry

Single-nucleated cells were isolated from perfused whole mice brains using Percoll™ gradient (Cytiva, cat. 17089101). Cells were analyzed with the following antibodies using either a myeloid panel (anti-CD45 APC-Cy7, BioLegend, cat.103116; anti-CD19 BUV786, Invitrogen, cat. 417-0193-82; anti-CD11b BV510, BioLegend cat. 101245; anti-CD11c BUV615, Invitrogen, cat. 366-0114-82; anti-Ly6C FITC, BioLegend, cat. 128005; anti-Ly6G BV421, BD Biosciences, cat. 560603; anti-CD163 PE, BioLegend, cat. 156703; anti-MHC-II(IE-IA) APC, Invitrogen, cat. 17-5321-82 and anti-CD68 PE-Cy7, Invitrogen, cat. 25-0681-82) or a lymphoid panel (anti-CD45 APC-Cy7, BioLegend, cat.103116; anti-CD11b BV510, BioLegend cat. 101245; anti-CD3 PE, BioLegend, cat. 100206; anti-CD4 PerCP, Invitrogen, cat. 45-0042-82; anti-CD8α APC, BioLegend, cat. 100712; anti-CD44 PE-Cy7, BioLegend, cat. 103030; anti-PD-1 BUV737, Invitrogen, cat. 367-9985-82; anti-FoxP3 BV421, BioLegend, cat. 126419 and anti-TCRγδ FITC, BioLegend, cat. 118105). Briefly, cells were suspended in 50 μl FACS buffer (0.5% BSA in PBS, EDTA 2mM) for cell surface marker analysis by staining with the appropriate antibodies for 30 min at 4 °C. Intracellular staining was achieved following manufacturer’s recommendations for the eBioscience™ Foxp3 / Transcription Factor Staining Buffer Set (Thermo Fisher Scientific, cat. 00-5523-00). Samples were pelleted and Countbright Beads (Thermo Fisher Scientific, cat. C36950) were added before analyzing the samples in a BD FACSymphony™ A5 SE flow cytometer.

Analysis was performed using FlowJo software. Gating strategy for the myeloid panel was performed with SSC-H / SSC-A□>□FSC-H / FSC-A□> SSC-A / FSC-A followed by fluorescence minus one (FMO), for CD45^+^, CD19^+^, Ly6G^+^, Ly6C^+^, CD68^+^ and CD163^+^ populations (**Supplementary Figure 1A**). Gating strategy for the lymphoid panel was performed with SSC-H/SSC-A□>□FSC-H/FSC-A□>□SSC-A/FSC-A followed by fluorescence minus one (FMO), for CD45^+^, CD3^+^, TCRγδ^+^, CD8^+^, CD4^+^ and FoxP3^+^ populations (**Supplementary Figure 1B**).

### 2.8. Lymphocyte restimulation *in vitro* and cytokine measurement

Cells were plated in 96 well plate in RPMI 1640 supplemented medium and restimulated with anti-mouse CD3ε clone 145-2C11(BioXCell, cat. BE0001-1) at 10μg/mL and MOG35-55 (GeneMed Synthesis, Inc.; sequence: MEVGWYRSPFSRVVHLYRNGK) at 20μg/mL for 72 hours. Supernatants were collected, flash frozen and kept at −80°C until analysis. LEGENDplex™ MU Th Cytokine Panel (12-plex) w/ VbP V03 (BioLegend, cat. 741044) was used to determine the concentrations of IFN-γ, IL-5, TNF-α, IL-2, IL-6, IL-4, IL-10, IL-9, IL-17A, IL17F, IL-22 and IL-13. The LEGENDplexTM Mouse Th Cytokine Panel is a bead-based multiplex assay panel, using fluorescence–encoded beads suitable for use on various flow cytometers. Samples were thawed extemporaneously, centrifuged and used undiluted. The assay was performed following the manufacturer’ s recommendations. Samples were analyzed on a BD FACSymphony™ A5 SE flow cytometer. Results were analyzed using the LEGENDplex™ Data Analysis Software Suite from VigeneTech. Concentrations were determined using a standard curve and were then normalized on the number of CD45+ cells per well.

### 2.9. Single cell RNAseq

#### Sample preparation

Single-cell libraries were generated using the Chromium GEM-X Single Cell 3′Reagent Kits v4 with Feature Barcode technology for Cell Surface Protein (10x Genomics, PN-1000691) following the manufacturer’s protocol. Single-cell suspensions were obtained from 16 mice (4 per experimental condition: unstressed male, stressed male, unstressed female, stressed female) and labeled with sample-specific barcoded antibodies (hashtag oligonucleotides) to enable multiplexing. CD45⁺ immune cells isolated from mouse brain were prepared and a total of 28,990 cells per condition were loaded into one lane of a GEM-X 3’ Chip, targeting an expected recovery of approximately 20,000 cells. Gel Bead-in-Emulsions (GEMs) were generated using the Chromium X (Wang Lab, Biological Sciences Platform of Sunnybrook Research Institute), followed by reverse transcription and cDNA amplification. For each reaction, 25% of total amplified cDNA was used for library construction according to the manufacturer’s instructions. Equal molar amounts of gene expression and cell surface protein libraries were pooled and sequenced on the Illumina NovaSeq X 10B flow cell platform (The Centre for Applied Genomics, The Hospital for Sick Children, Toronto).

#### Single-cell RNA-seq analysis

Demultiplexing, alignment, barcode extraction, and UMI-collapsing were performed using the Cell Ranger Pipeline (v7.2.0, 10x Genomics) with the mouse reference genome. Downstream single-cell transcriptomic analyses were conducted by pooling cells from all four mice per experimental condition using R (v4.4.0) and the Seurat package (v5.2.1).

We removed low-quality cells and potential red blood cell contaminants determined by abnormal unique number of genes (>6000 or <200), high mitochondria content (>10%) and high hemoglobin gene expression (>1%). Doublets were identified and removed using scDblFinder prior to normalization with SCTransform. Elbow plot were used to select distinguished principal components, and the top 30 PCs were included for downstream analysis. Samples were then integrated for batch correction using the Harmony algorithm. The cells were clustered via FindNeighbors and FindClusters functions, setting resolution to 0.5 using the reduction result from harmony. Uniform Manifold Approximation and Projection (UMAP) was used to visualize the results.

Major lymphocyte lineages were annotated using canonical marker genes, and differentially expressed genes (DEGs) were identified using FindAllMarkers for cluster markers and FindMarkers for pairwise group comparisons. Tissue-resident lymphocyte clusters were defined based on the combined gene expression. Module scores were computed using AddModuleScore for specific gene signatures, including Th1/Th2/Th17 cytokine response, and transcriptional regulatory modules. Condition comparisons (e.g., UCMS vs unstressed and female vs male) were performed using Wilcoxon rank-sum tests, and adjusted p-values were computed using the Benjamini–Hochberg correction. For visualization, DotPlots were used to display average expression and fraction of expressing cells for lineage, activation, Th1/Th2/Th17, co-stimulatory, and chemokine receptor gene sets. Heatmaps were generated using pheatmap and ggplot2, based on averaged expression matrices or module score statistics; where indicated, expression values were row-wise Z-score normalized to visualize relative expression patterns across genes. Volcano plots were created using ggplot2, highlighting significantly up- and downregulated genes (adjusted p < 0.05, |log₂FC| > log₂(1.5)).

### 2.10. Statistical Analysis

GraphPad Prism 10 software was used for the analyses. Normality was assessed using the Shapiro–Wilk and Kolmogorov–Smirnov tests and equality of variances was verified using the F-test of equality of variance. The significance threshold used was p < 0.05. Specific statistical tests are described in figure legends.

## 3. Results

### Unpredictable Chronic Mild Stress (UCMS) promotes depressive-like behavior

To investigate the lymphocyte landscape in the chronically stressed brain, we first validated the UCMS model in adult age-matched female and male C57BL/6 mice assessing changes in weight, serum corticosterone levels and depressive-like or related behaviors (**Figure 1A**). As expected, both female and male mice have significantly reduced weight gain and increased serum corticosterone when subjected to UCMS (**Figure 1B, C**). Next, we evaluated depressive-like behaviors of mice with 0-, 4-, or 8-weeks (wk) of UCMS (**Figure 1D** and **Supplementary Figure 1**). In both sexes, UCMS exposure led to increased time spent in the outer zone of an Open Field Test (OFT), indicative of heightened anxiety, and increased time spent immobile in the Forced Swimming Test (FST), indicative of enhanced passive coping which is a core feature of depression. Interestingly, female but not male mice exhibited reduced social interaction in the Social Interaction Test (SIT) (**Supplementary Figure 1**) and decreased preference for sucrose water in the Sucrose Preference Test (SPT), suggesting blunted reward sensitivity and the emergence of anhedonia. This finding aligns with previous studies using alternative depression models (Page et al. 2016) and with clinical observations that anhedonia is more prevalent in women than in men (Albert 2015; Young 1998). Thus, the UCMS model effectively induces depressive-like phenotypes in both sexes while also uncovering sex-specific differences consistent with clinical observations.

**Figure 1.**
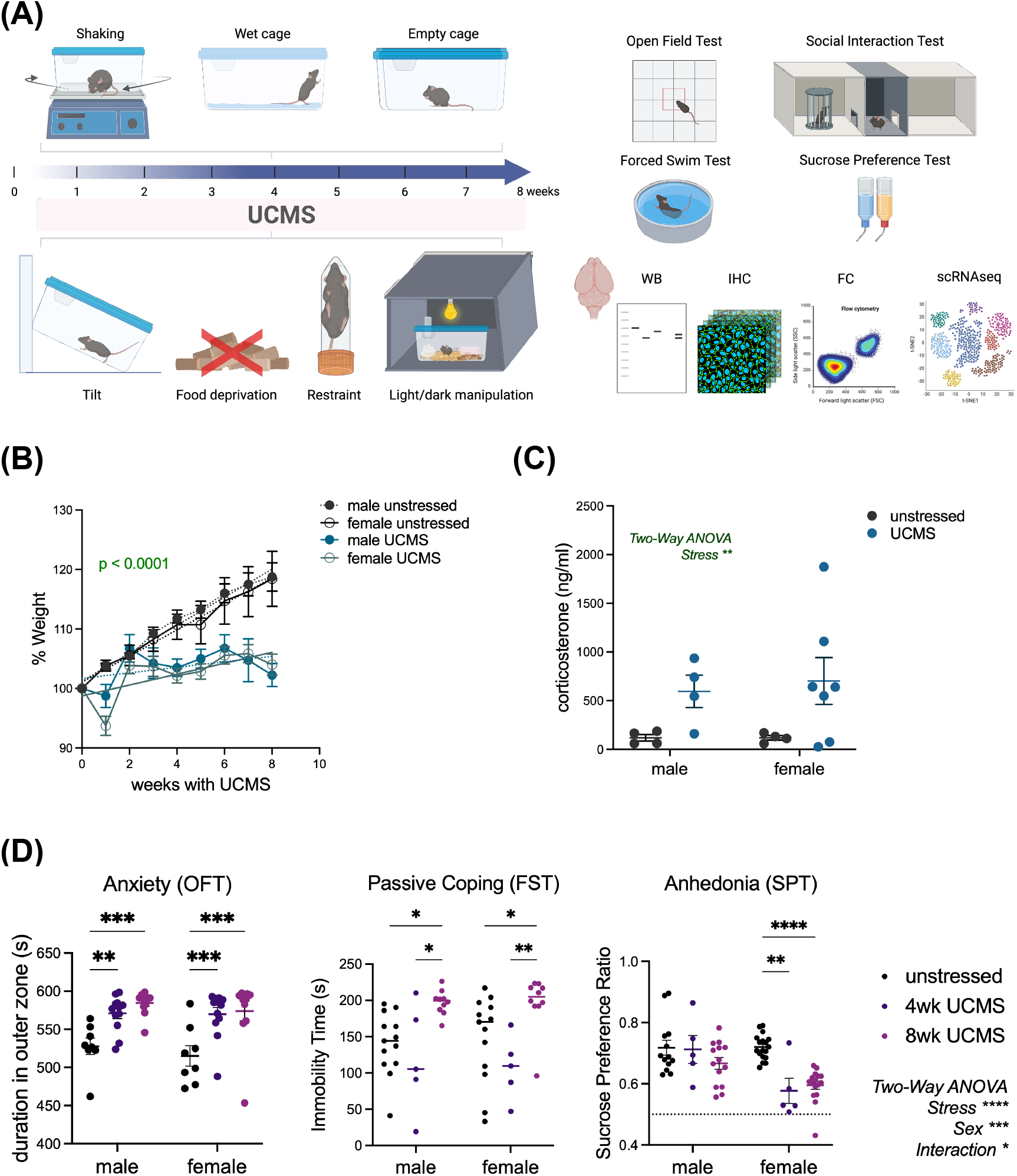
UCMS induces depressive-like behavior in female and male mice. **(A)** Eight-week-old mice were either left unstressed or subjected to an 8-week UCMS. Changes in %weight (**B)**, serum corticosterone level **(C)** and depressive-like behavioral **(D)** are reported where each dot is a mouse. A was analyzed by linear regression. B and C were analyzed by two-way ANOVA with Tukey’s multiple comparisons test. D was analyzed with student t test. * p < 0.05; ** p < 0.01; *** p < 0.001; **** p < 0.0001.

### UCMS alters hippocampal neuron organization

To analyze the general neuronal health of the hippocampus we assessed the protein expression levels of markers of synaptic plasticity: Synaptophysin (SYN) and Post-Synaptic Density protein 95 (PSD95), which are pre- and post-synaptic vesicles proteins respectively. We observed a reduction of PSD95 expression in female, but not male, hippocampi under chronic stress conditions (1.37 fold decrease, p < 0.05) (**Figure 2A**). While there were no variations in the expression levels of synaptophysin following UCMS protocol, both proteins showed a significant sex bias with significantly higher levels expressed in males compared to females (2way ANOVA sex variation p <0.0001, **Figure 2A**).

**Figure 2.**
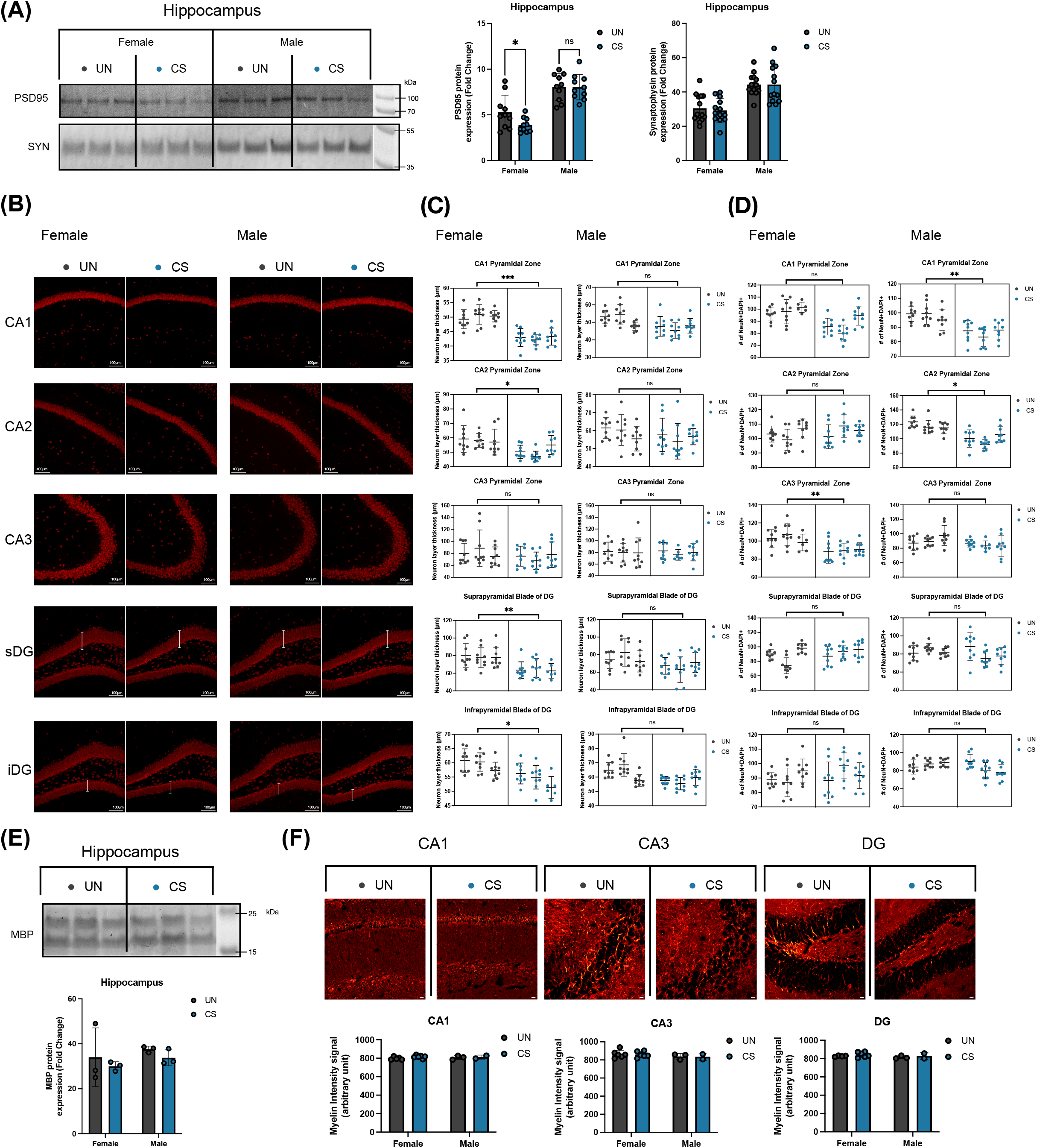
UCMS alters hippocampal neuron organization but not myelin integrity. Eight-week-old C57BL/6 mice were either left unstressed (UN) or subjected to 8-wk UCMS (CS). Hippocampi were processed either for protein extraction or for immunohistochemistry. **(A)** Western Blot showing PSD95 and Synpatophysin (SYN) with their protein expression normalized using Total Protein Quantification. Data are shown as mean ± SD. Two-way ANOVA with Fisher’s LSD multiple comparisons test was used. Each replicate corresponds to an individual mouse. **(B)** Representative images of Red, NeuN expression in the CA1, CA2, CA3 pyramidal and dentate gyrus (DG; subdivided into suprapyramidal and infrapyramidal blades) regions of the hippocampus in male and female. **(C)** Quantification of neuronal thickness, data is presented as mean ± SD and analyzed using nested t-test. **(D)** Quantification of NeuN+ DAPI+ double positive cells, data is presented as mean ± SD and analyzed using nested t-test. **(E)** Western Blot showing Myelin-Basic Protein (MBP) expression in female mice and its protein expression normalized using Total Protein Quantification. **(F)** SRS imaging of CH2 at 2845 cm-1 of the hippocampus from CA1 region, scale bar: 20□μm; CA3 region, scale bar: 20□μm; and Dentate Gyrus, scale bar: 10□μm. Adjunct histograms of the the myelin signal intensity quantification.

The Cornu Ammonis CA1, CA2 and CA3 and Dentate Gyrus (DG) regions of the hippocampus are crucial for memory, learning and social behavior. Previously, alterations of hippocampal neurons have been reported under chronic stress. Here we report a reduction in the thickness of the neuron layers of 15.9% in the CA1 pyramidal zone (Δ of thickness = −7.38 ± 1.28 μm, p < 0.001), 13.5% in the CA2 pyramidal zone (Δ of thickness = −7.33 ± 4.70 μm, p < 0.05) but not CA3, and both infrapyramidal (9.2% reduction, Δ of thickness = −5.25 ± 0.71 μm, p < 0.05) and suprapyramidal (20.3% reduction, Δ of thickness = −14.45 ± 2.63 μm, p < 0.01) blades of the DG in female hippocampi following UCMS protocol (**Figure 2B, C**). These differences were abscent from the male hippocampi after chronic stress exposure. The pyramidal neuron layer thickness was also comparable between males and females. To better characterize the changes seen in these regions, we counted the number of neurons by immunohistochemistry (IHC). The reduction in thickness of the CA2 and DG layers of neurons in female hippocampi under chronic stress was not associated with reduction in the number of neurons (**Figure 2D**). Thus, the size reduction of these specific regions cannot be characterized as an atrophy but rather suggests an alteration of the density of the neurons and possibly of their morphology. Neuron counts were found reduced in the CA1 (12.7% decrease, NS p=0.0625) and CA3 (13.9% decrease, p < 0.01) pyramidal zone of females after UCMS protocol (**Figure 2D**). Male hippocampal neuron counts were reduced in both CA1 and CA2 regions (12.6% reduction, p < 0.01 and 17.2% reduction, p < 0.05 respectively) but not in CA3 and DG zones (**Figure 2D**).

Overall, female mice exposed to the UCMS protocol exhibit more pronounced alterations in hippocampal neuronal organization, characterized by reduced density in the CA1 region whereas male counterparts displayed a milder phenotype, marked primarily by a decrease in neuronal numbers without evident histological atrophy in the same region.

### UCMS does not affect the hippocampal myelination profile

To assess myelin integrity and potential abnormalities in the hippocampus we first measured the expression levels of Myelin-Basic Protein (MBP) by western blot. We observed no difference in the expression of MBP in the hippocampus of both male and female mice exposed to the UCMS protocol when compared to controls (**Figure 2A**). Next, we used stimulated Raman scattering (SRS) microscopy on brain slices to specifically quantify myelin content across several regions of interest in the hippocampus, namely CA1, CA3 and the Dentate Gyrus. Raman spectroscopy leverages quantitative detection of endogenous metabolites in a label-free manner by assessing the vibrations of chemical bonds. Raman scattering microscopy combines this technique with targeted fingerprints of predetermined molecules and offers subcellular resolution. Similarly to MBP expression levels, we did not report any changes in the myelin content and integrity in these regions when comparing sex or chronic stress status (**Figure 2B**). Overall, we documented no changes in the myelination profile of the hippocampus of the mice following the UCMS protocol.

### Preferential activation of neutrophils and CD4 T cells in male and CD8 T cells in female UCMS brain

To evaluate the type of immune cells in the UCMS brain, we optimized two flow cytometry panels and analyzed all CD45^high^ hematopoietic cells (**Supplementary Figure 3** and **Methods**). We identified distinct CD11b^+^ myeloid (Ly6G^+^ Neutrophils, CD68^+^ Macrophages, CD163^+^CD68^+^ Border Associated Macrophages-BAM, and Ly6C^+^ myeloid cells) and CD11b^-^lymphocytes (CD19^+^ B cells, CD3^+^TCRγδ^+^ γδT cells, conventional CD3^+^CD4^+^ T cells, CD3^+^CD8^+^ T cells, and CD4^+^Foxp3^+^ regulatory T cells-Tregs). Strikingly, the types of immune cells in the female and male brains differ significantly. The female brain has a higher number of macrophages and BAM, whereas the male brain has a higher number of neutrophils, Ly6C^+^ myeloid cells, CD4 T cells and Tregs (**Figure 3A**). Importantly, no significant change was found in the number of immune cell populations comparing unstressed (UN) and UCMS (CS) mice except neutrophils (**Figure 3B**). Male, but not female, UCMS mice have reduced numbers of neutrophils in the brain of UCMS mice (**Figure 3B**).

**Figure 3.**
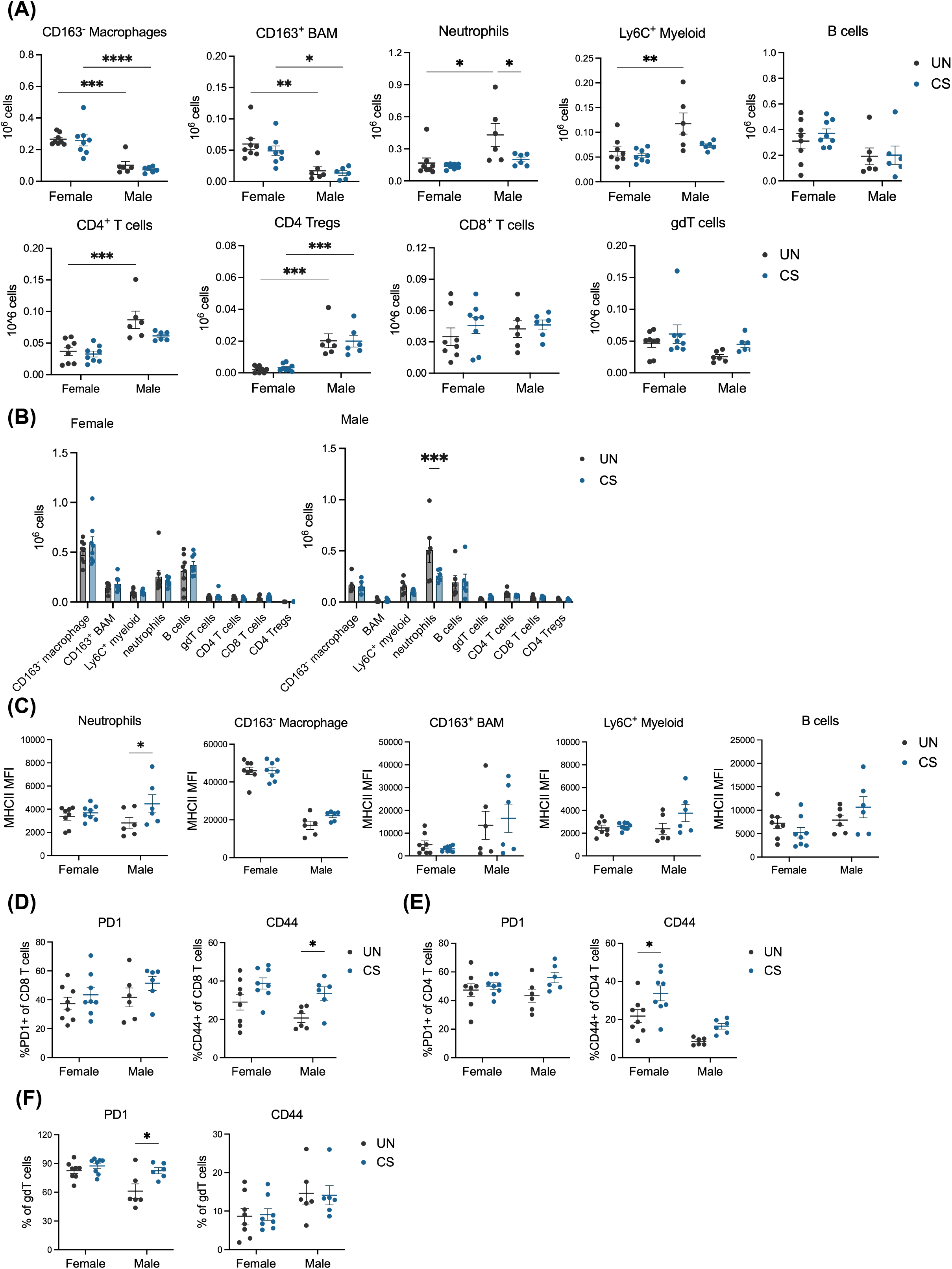
Preferential activation of neutrophils and CD4 T cells in male and CD8 T cells in female UCMS brain. Eight-week-old C57BL/6 mice were either left unstressed or subjected to 8-wk UCMS. Whole brains were dissected, enzymatically digested and cells isolated by percoll gradient followed by flow cytometry analysis. **(A)** number of different immune cell population per brain; **(B)** MHC class II expression (Median Fluorescence intensity, MFI) are reported. C, D. PD1 and CD44 expression are ploted for CD4 **(C)** and CD8 **(D)** T cells. Each dot is a mouse in all figures. Two-way ANOVA with Tukey’s multiple comparisons test was used. * p < 0.05; *** p < 0.001.

Next, we analyzed the activation status of immune cells. UCMS induced a significant increase in the expression of MHC class II (MHCII) on neutrophils in male but not female mice (**Figure 3C**). MHCII expression was not different on other myeloid populations or B cells (**Figure 3C**). Interestingly, male UCMS mice also have a higher frequency of CD8 T cells that expressed CD44 but not PD1, suggesting increased activation that is likely independent of CD4 T cell help (**Figure 3D**). In contrast, female mice have more activated CD4 T cells (**Figure 3E**). We did not find a difference in PD1 or CD44 expression in γδT cells (**Figure 3F**). Thus, while the overall number of different immune cell populations in the brain does not change in UCMS, there are selective activation of lymphocytes that may be sex-dependent.

### Brain tissue resident T, NKT and γδT cells have distinct type 1, 2 and 17 transcriptomes

Given the limited resolution of flow cytometry in distinguishing lymphocyte lineages, we use single-cell RNA sequencing (scRNAseq) to comprehensively profile the immune cell landscape and identify the cellular sources of T helper-like cytokines in the UCMS brain. This analysis examined lymphocyte composition and cell state in 8 female and 8 male mice that were unstressed or subjected to the UCMS protocol. After quality control and batch correction (**Methods**), 6720 number of lymphocytes were analyzed and visualized in Uniform Manifold Approximation and Projection (UMAP), constituting of 19 clusters (**Figure 4A**). We identified major lymphocyte lineages using established markers (**Figure 4B**): αβT cells (Cluster 4, 5, 6 and 10: *Trac, Trbc, CD3d, Cd3g, CD247*), γδT cells (Cluster 9, 12 and 17: *Trdc, Trgc, Trgv1, Trgv6, Zbtb16*), NK cells (Cluster 3: *Klrb1c* (NK1.1)*, Nkg7, Ncr1* (Nkp46)*, Klrk1* (NKG2D)*, Klrc1* (NKG2A)*, Klrd1* (CD94)*, Klra7, Klra8*), NKT cells (Cluster 14: *Trac, Trbc, Cd3d, Cd3g, Cd247, Klrb1c, Cd1d*) and B cells (Clusters 0, 1, 2, 8, 11, 13 and 16: *CD19, Cd79a, Ms4a1, Blnk, Ighd*).

**Figure 4.**
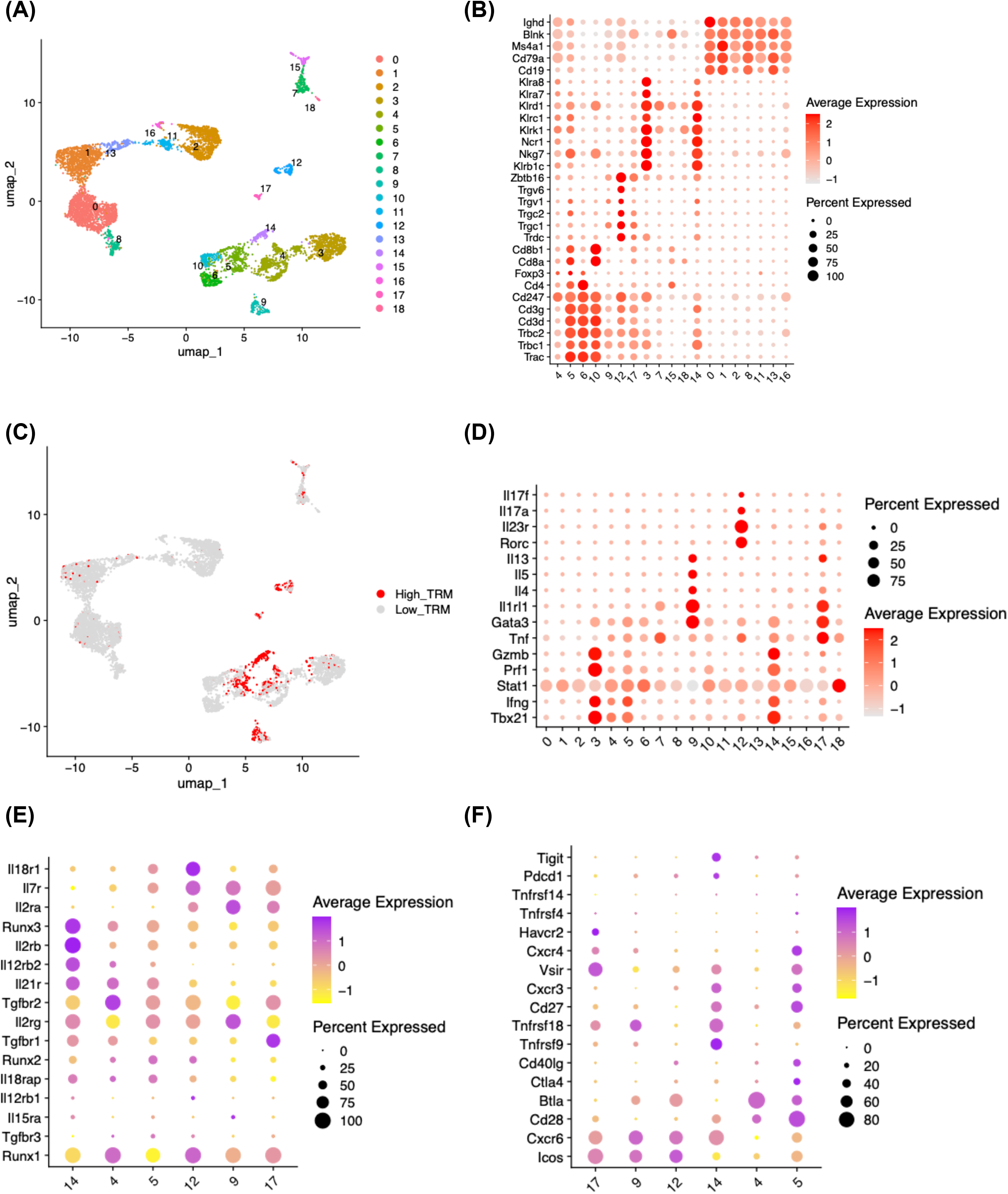
Brain tissue resident T, NKT and γδT cells have distinct type 1, 2 and 17 transcriptomes with selective regulatory network. **(A)** UMAP visualization of lymphocyte revealing 19 transcriptionally distinct subclusters. **(B)** Dot plot showing the expression of canonical markers defining major lymphocyte lineages, including αβ T, γδ T, NK, NKT and B cells. **(C)** UMAP highlighting tissue-resident lymphocytes (TRM), where cells within the top 10% of the TRM module score (Δ = TRM_Pos − TRM_Neg) are shown in red. **(D)** Dot plot illustrating representative signature genes of Type 1, Type 2, and Type 17 lymphocytes. **(E-F)** Clustered dot plots showing expression profiles of cytokine/chemokine receptors, transcription factors, and co-stimulatory/co-inhibitory molecules across tissue resident lymphocytes clusters. Genes and clusters were hierarchically ordered by similarity in average expression.

To assess whether the brain lymphocytes are tissue resident or circulatory, we analyzed genes involved in tissue residency (Christo et al. 2024) including *Cd69*, *Itgae* (CD103), *Itga1* (Ly49a), *Cxcr6*, *Zfp683*, *Prdm1* and negative regulators *S1pr1, Klf2* and *Ccr7* (**Figure 4C** and **Supplementary Figure 4A**). We identified several clusters enriched for tissue-resident lymphocyte markers, including NKT cells (clusters 14), αβT cells (clusters 4 and 5) and all γδT cells (clusters 9, 12 and 17). Whereas the remaining αβT cell cluster (cluster 6 and 10) exhibited a naïve-like phenotype corresponding to CD4 and CD8 T cells, respectively, enriched with naïve markers such as *Tcf7, Lef1* and negative for activation markers such as *Cd44 and Slamf7* (**Supplementary Figure 4B**). Thus, several brain lymphocyte clusters were identified that are likely tissue resident and are of distinct T cell lineages including NKT, γδT and αβT cells.

Lastly, we directly analyzed the expression of type 1, type 2, and type 17 signature genes (**Figure 4D**). Strikingly, a γδT cell cluster (rγδT17, Cluster 12) is enriched for type 17 signature genes *Rorc, Il23r, Il17a, Il17f* and the remaining γδT cells (rγδT2, clusters 9 and 17) uniquely express type 2 signature genes including *Gata3*, *Il1rl1* (ST2), *Il4, Il5, Il13*. On the other hand, tissue resident αβT cells (rTc1, clusters 4 and 5) and NKT cells (rNKT1, clusters 14) are all enriched for type 1 signature genes including *Tbx21* (TBET) and *Ifng*, similarly to NK cells (cluster 3) which also have high expression of *Prf1* (Perforin) and *Gzmb* (Granzyme B). Of note, cluster 4 and 5 contains both CD4^+^ and CD8^+^ αβ T cells (**Figure 4B**), hereafter referred as rTc1 (“c” for cytotoxicity) for simplicity.

Thus, the different types of T helper-like responses observed in the brain correspond to specific lymphocyte lineages that are enriched for tissue-resident markers. This finding is consistent with prior findings that tissue-resident lymphocytes are potent cytokine. Accordingly, we focused our subsequent analyses on these cells.

### Selective expression of Runx family transcription factor, cytokine receptors, and checkpoint molecules in brain tissue resident lymphocytes

The observation of distinct differentiation states among brain resident lymphocytes prompted us to analyze potential regulators. Previous studies have shown that tissue residency is maintained by IL-15 and TGF-β signaling and can be modulated by IL-12 and IL-18 (Christo et al. 2024). Accordingly, we analyzed the transcript expression of key cytokine receptors and transcription factors associated with these pathways (**Figure 4E** and **Supplementary Table 1**). Overall, comparing to the αβT and NKT cells, the tissue resident γδT cell clusters have higher expression of receptors to IL-18, IL-15, IL-12, IL-7, IL-2, and TGF-β. Notably, rγδT17 cells have the highest expression of *Il18r1* (*adj p* = 2.09E-114, LOG_2_ fold = 3.35) and *Il12rb1* (*adj p* = 1.12E-29, lOG_2_ fold = 3.39) suggesting a higher potential responsiveness to IL-12/IL-23 and IL-18, drivers of pathogenic type 1 and 17 responses (Tang et al. 2025; Pawlak et al. 2022). Conversely, the rγδT2 cluster 9 has the highest expression of receptors for IL-2 (*Il2ra*, *adj p* = 0, LOG_2_ fold = 5.11; *Il2rg*, *adj p* = 9.68E-12, LOG_2_ fold = 0.73) and IL-15 (*Il15ra, adj p* = 4.87E-11, LOG_2_ fold = 2.42); whereas the rγδT2 cluster 17 has the highest expression of TGF-β receptor (*Tgfbr1*, *adj p* = 9.99E-8, LOG_2_ fold = 1.71). In contrast, rNKT1 preferentially expresses receptor to IL-12 and IL-21 (*Il21r*, *adj p* = 1.67E-14, LOG_2_ fold = 1.18; and *Il12rb2*, *adj p* = 4.47E-85, LOG_2_ fold = 1.83) and rTc1 cluster 5 preferentially expresses TGF-β receptor 3 (*Tgfbr3, adj p* = 6.61E-23, LOG_2_ fold = 1.02).

Runx family transcription factors were previously shown to be essential for tissue residency of lymphocytes (Fonseca et al. 2022; Milner et al. 2017). We found significantly higher levels of *Runx1* and *Runx 2* in rγδT17 and rTc1 cells (rγδT17: *Runx1, adj p* = 6.04E-12, LOG_2_ fold = 0.7; *Runx2*, *adj p* = 5.16E-8, LOG_2_ fold = 0.95; rTc1 cluster 4: *Runx1* cluster 4, *adj p* = 1.32E-26, LOG_2_ fold = 0.7; *Runx2*, *adj p* = 1.14E-11, LOG_2_ fold = 0.92; rTc1 cluster 5, *Runx2*, *adj p* = 8.18E-54, LOG_2_ fold = 0.98). Lastly, *Runx3* is preferentially enriched in rNKT1 cells (*adj p* = 7.67E-55, LOG_2_ fold = 2.38).

Next, we analyzed the expression of costimulatory, coinhibitory molecules and chemokine receptors to further understand the regulation of tissue resident lymphocytes (**Figure 4F** and **Supplementary Table 1**). Comparing to αβT and NKT cells, the rγδT cells preferentially express *Icos* (clusters 9, *adj p* = 4.21E-89, LOG_2_ fold = 3.14; cluster 12, *adj p* = 7.96E-84, LOG_2_ fold = 3.54; and cluster 17, *adj p* = 6.29E-40, LOG_2_ fold = 2.65) and *Cxcr6* (clusters 9, *adj p* = 1.74E-162, LOG_2_ fold = 3.72; cluster 12, *adj p* = 5.19E-124, LOG_2_ fold = 3.38; and cluster 17, *adj p* = 1.95E-39, LOG_2_ fold = 2.53). Of the rγδT clusters, rγδT2 clusters have higher expression of Vista (*Vsir, cluster 17, adj p = 2.09E-15,* LOG_2_ fold = 1.53), GITR (*Tnfrsf18,* cluster 9*, adj p* = 7.57E-78, LOG_2_ fold = 3.13; cluster 17, *adj p* = 7.61E-12, LOG_2_ fold = 2.08) and Tim3 (*Havcr2,* cluster 17, *adj p* = 3.45E-32, LOG_2_ fold = 3.09). In contrast to γδT cells, rTc1, most notably cluster 5, exhibit elevated expression of the B7 family members *Cd28* (clusters 4, *adj p* = 2.08E-65, LOG_2_ fold = 2.16; and cluster 5, *adj p* = 1.86E-261, LOG_2_ fold = 2.66) and *Ctla4* (cluster 5, *adj p* = 2.23E-90, LOG_2_ fold = 3.35) and the chemokine receptors *Cxcr3* (*adj p* = 4.4E-269, LOG_2_ fold = 4.19) and *Cxcr4* (clusters 5, *adj p* = 2.68E-6, LOG_2_ fold = 0.45). The rTc1 cluster 5 also have higher expression of other costimulatory/coinhibitory molecules such as *Cd27* and *Vsir*. Conversely, rNKT1 cells (cluster 14) are uniquely enriched for *Tnfrsf9* (4-1BB, *adj p* = 1.98E-110, LOG_2_ fold = 3.32), *Pdcd1* (PD-1, *adj p* = 5.64E-50, LOG_2_ fold = 3.57) and *Tigit (adj p* = 1.49E-108, LOG_2_ fold = 3.7). Finally, both rγδT17 and rTc1 cells significantly increased *Cd40lg* (CD40) expression (cluster 4, *adj p* = 1.86E-10, LOG_2_ fold = 1.82; cluster 5, *adj p* = 2.27E-129, LOG_2_ fold = 3.16; cluster 12, *adj p* = 6.91E-5, LOG_2_ fold = 2.08), alluding potential ability to provide T cell help to myeloid cells.

Thus, distinct cytokine receptor, transcription factor and costimulatory/coinhibitory molecules are associated with different subsets of resident Tc1, NKT1, γδT2, and γδT17 cells. The difference in the gene regulatory network is likely in response to differences in their microenvironment.

### Chronic stress expands tissue resident lymphocytes with sex-divergent flavors

To investigate whether sex or chronic stress alters the T helper-like responses in the brain, we analyzed the cluster frequencies and the enrichment of type 1, 2, and 17 signatures in tissue resident lymphocyte clusters (**Figure 5A-D**). First, we analyzed the compositional changes of the various tissue resident clusters. We found that UCMS significantly expanded tissue resident clusters. These clusters are rTc1 (cluster 4), rγδT2 (cluster 9) and rNKT1 (cluster 14) in both sexes with a more prominent effect in females (**Figure 5A, B**). This finding likely reflects an increase in the number of tissue resident cells in response to UCMS in the brain as Figure 3B shows that there is no change in the overall numbers of hematopoietic cells in this context. On the other hand, sex had a more dominant effect on all αβT and γδT clusters: resident γδT cell clusters are significantly enriched in females whereas the resident Tc1 clusters are enriched in males in both unstressed and UCMS conditions with a more prominent sex bias in UCMS.

**Figure 5.**
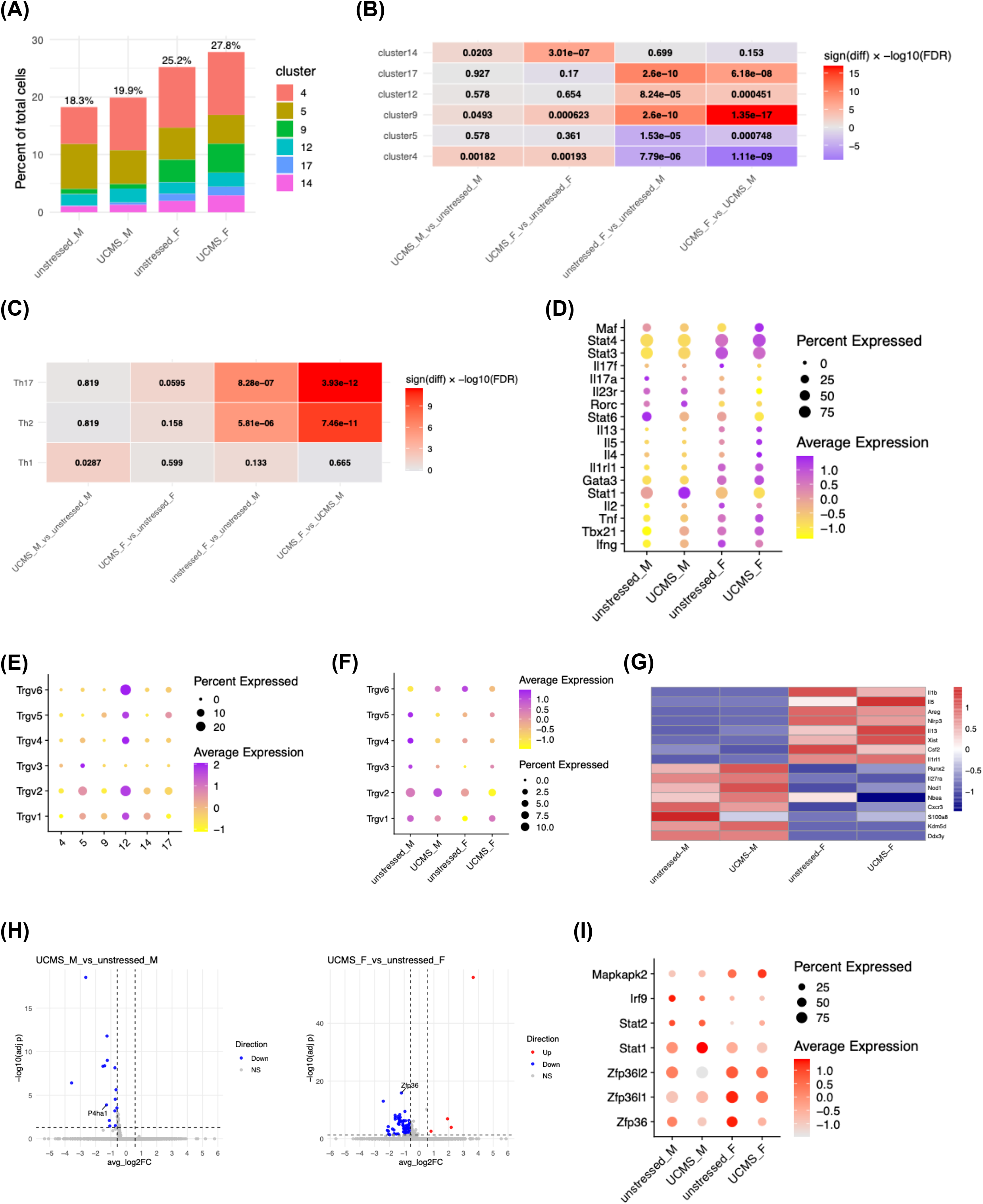
Chronic stress decreases *P4ha1* in male and *Zfp36* in female tissue resident lymphocytes, promoting type 1 and 17 responses respectively. **(A)** Percent of tissue-resident lymphocytes across sex and stress conditions, shown as stacked bars by cluster. **(B)** Heatmap of module-score differences (Wilcoxon test) for top10 marker genes of tissue-resident lymphocytes clusters across groups and annotated with adjusted p values. **(C-D)** Th1, Th2, and Th17-related module score, shown by comparison (Wilcoxon test) heatmap and expression of representative signature genes. **(E–F)** Expression of Trgv genes across tissue-resident lymphocytes clusters, and sex and stress conditions. **(G)** Heatmap of sex-differentially expressed genes **(F vs M)** across conditions. **(H)** Volcano plots of group contrasts highlighting significant genes. **(I)** Expression of translational repressors across tissue resident lymphocyte clusters.

Next, we compared the enrichment of type 1, 2 and 17 core signature genes across sex and stress conditions (**Figure 5C-D**). Notably, sex has a more dominant effect: female tissue resident lymphocytes have a significant enrichment of type 2 and 17 signature as compared to the male counterparts and that UCMS exacerbated this sex bias (**Figure 5C**). We also found that UCMS selectively increased type 1 signature genes in male but not female lymphocytes (**Figure 5C**, *adj p* = 0.0287). The enrichment in males is driven by type 1 signature genes such as *Tbx21, Ifng and Stat1* (**Figure 5D**). Instead, female tissue resident lymphocytes have a trend of type 17 signature enrichment with higher expression of *Stat4* and *Maf* but not downstream type 17 genes (**Figure 5D**), suggesting a nuisance effect.

### Stress-dependent γδT lymphocyte expansion is not associated with intestinal origin

Previous work suggested that stress-induced expansion of brain γδT17 cells in male mice were derived from the intestine in the chronic social stress model (Zhu et al. 2023). c-MAF promotes the differentiation of regulatory gut Th17 cells (Brockmann et al. 2023). To investigate whether the expansion of tissue resident lymphocytes in UCMS might similarly derive from the gut, we analyzed the variable γ chain regions (Vγ) of the T cell receptor (TCR) which defines distinct γδT substs. Intestinal intraepithelial γδT cells are mostly Vγ7^+^ (Trgv7), whereas brain meningeal γδT cells are predominantly Vγ6^+^ (Nielsen et al. 2017; Alves de Lima et al. 2020; Ribeiro et al. 2019). To this end, we did not detect any transcript of Vγ7 (*Trgv7*). Instead rγδT17 cells in our dataset were more enriched for *Trgv6* (Vγ6) and *Trgv2* (Vγ2) (**Figure 5E**), and more aligned with meningeal γδT17 cells. Next, we assessed the impact of stress and sex on γ variable chain usage. We found that *Trgv6* expanded in response to UCMS in male, but was more restricted in females. Instead, *Trgv1* and *Trgv3* expanded in females, alluding to sex-specific antigenic pressure during chronic stress (**Figure 5F**).

A subset of αβT cells, notably MAIT cells, express MR1, a non-classical MHC class I protein that presents microbial products (e.g. riboflavin-derivative antigens) to T cells. We found MR1 to be enriched rTc1 cluster 5 and not cluster 4 (**Supplementary Figure 5A-B**). However, unlike MAIT cells described in the mucosal sites which have restricted usage of T cell receptor alpha and beta chains (Godfrey et al. 2019), we found broad TCR alpha and beta chain usage in MR1^+^ cells (**Supplementary Figure 5A-B**).

Thus, our findings do not support an intestinal origin of either tissue-resident αβT or γδT cells within the UCMS brain.

### Chronic stress decreases *P4ha1* in male and *Zfp36* in female tissue resident lymphocytes

To investigate molecular regulators of the stress- and sex-driven effects on type 1, 2 and 17 responses, we analyzed differentially expressed genes (DEG, **Methods**). First, we analyzed sex-specific effects. We found 633 DEG (*adj p* < 0.05, |fold change| > 1.5) between male and female tissue resident lymphocytes in both unstressed and UCMS brain (**Figure 4G** and **Supplementary table 2**). As expected, X and Y chromosome linked genes are found to be differentially expressed such as *Xist* and *Ddx3y*. Notably, female lymphocytes have increased expression of *Nlrp3* and *Il1b*, genes related with inflammasome, as well as type 2 genes *Il5* and *Il13*, and *Csf2* which mediates myeloid cell recruitment. The higher Csf2 expression in females aligns with the increased abundance of brain macrophages observed by flow cytometry (**Figure 3A**). In contrast, male lymphocytes exhibit higher expression of *Il27ra* and *Cxcr3*, cytokine and chemokine receptors that are associated with type 1 responses, as well as *Nod1*, an intracellular pattern recognition receptor known to recognize and clear microbial infections (**Figure 4G**).

Next, we analyzed stress-driven effects. We identified 91 DEG in females and 15 DEG (*adj p* < 0.05, |fold change| > 1.5) in males comparing UCMS and unstressed controls (**Figure 5H-I** and **Supplementary Table 3**). We found that tissue resident lymphocytes in UCMS brain significantly reduced the transcript expression of *Zfp36* (Tristetraprolin, TTP) in females and *P4ha1* (prolyl 4-hydroxylase 1) in males (**Figure 5H**). P4HA1 suppressed IFNγ and TNFα production from CD8 T cells (Ma et al. 2025). TTP is an RNA binding protein that promotes the post-transcriptional degradation of mRNA of various inflammatory cytokines (Fu and Blackshear 2017; Makita et al. 2021). Previous work show that TTP in T cells restrains IL-17 production in aged mice (Peng et al. 2020). TTP activity can be regulated by MAPK activated protein kinase 2 (Mk2) and is dependent on phosphorylation (Stoecklin et al. 2004). Transcriptionally, TTP is regulated by interferon-induced gene factor IGSF3 (Yadav et al. 2024), a transcription factor complex formed by STAT2, IRF9 and when activated engages with STAT1 (Platanitis et al. 2019). We analyzed the expression of these genes along with two other ZFP36 family members *Zfp36l1* and *Zfp36l2* across tissue resident lymphocyte clusters (**Figure 5I** and **supplementary Table 1**). We found that both Zfp36 and Zfp36l1 are more enriched in γδT2 cells (zfp36, cluster 17: *adj p* = 5.01E-18, LOG_2_ fold = 2.04; zfp36l1, cluster 9, *adj p* = 9.39E-19, LOG_2_ fold = 1.21) whereas Zfp36l2 is more enriched in rTc1 (cluster 5, *adj p* = 2.2E-27, LOG_2_ fold = 0.86) and rNKT1 cells (*adj p* = 1.14E-19, LOG_2_ Fold = 1.36) (**Figure 5I** and **supplementary Table 1**). γδT2 cells are also enriched for Mk2 expression (*Mapkapk2, adj p* = 8.28E-12, LOG_2_ fold = 1.76) whereas only Tc1 cells (cluster 5) are enriched for *Stat1* expression (*adj p* = 2.49E-6, LOG_2_ fold = 0.55).

Thus, our analysis support sex-specific differences in tissue-resident lymphocytes and suggests that UCMS may potentiate lymphocyte activities by suppressing negative regulators of cytokine responses.

### Sex-specific T lymphocyte responses to chronic stress

Both P4HA1 and ZFP36 are suppressors of cytokine production. As chronic stress decreased the expression of P4HA1 and ZFP36 in male and female brain resident lymphocytes, respectively, we tested the hypothesis that stressed brain lymphocytes have exacerbated cytokine production. To this end, we isolated immune cells from the brains of unstressed and UCMS mice and restimulated them *in vitro*. Stimulation with anti-CD3 was used to specifically activate the CD3-TCR complex on αβT cells and γδT cells. MOG peptide 35-55 was used as a negative control for antigen-specific stimulation as it is the main auto-antigen characterized in autoimmune T cell mediated demyelination studies. Twelve cytokines were analyzed and grouped into type 1, 2 and 17 signatures. Consistent with the scRNAseq finding, Type 1 cytokines, IFN-γ, IL-2 and TNF-α were found significantly elevated in brain lymphocytes from male mice that underwent the UCMS protocol (5.1 fold for IFN-γ, p < 0.05; 3.4 fold for IL-2, p < 0.05 and 2.4 fold for TNF-α, p < 0.01 respectively) but remained unchanged in females or decreased in the case of IL-2 (2.5 fold, p < 0.001) (**Figure 6A**). Cytokines associated with type 2 signature showed no secretion of IL-4 and IL-13 and very little secretion of IL-5 although higher in females compared to males and significantly increased in chronic stress in both sexes (**Figure 6B**). Female brain lymphocytes showed overall higher levels of IL-17A and IL-17F than males with both cytokines further elevated in chronic stress conditions (2 fold for IL-17A, p < 0.05 and 2.3 fold for IL-17F, p < 0.05) (**Figure 6C**). IL-9 production was minimal (Means < 5pg/mL) and IL-22 concentration was decreased in brain lymphocytes from chronically stressed female mice (1.2 fold, p < 0.05). While male lymphocytes exhibited a similar increasing trend of IL-17A (2.2 fold, p=0.114) and IL-17F (2.4 fold, p < 0.05) increase in chronic stress, they also display elevated levels of IL-9 (3.3 fold, p < 0.05) and IL-22 (9.9 fold, p < 0.05) which are not seen in females (**Figure 6C**). Finally IL-6 and IL-10 remained unchanged across both sexes and stress conditions, with notably high levels of IL-6 and low levels of IL-10 (**Figure 6D**).

**Figure 6.**
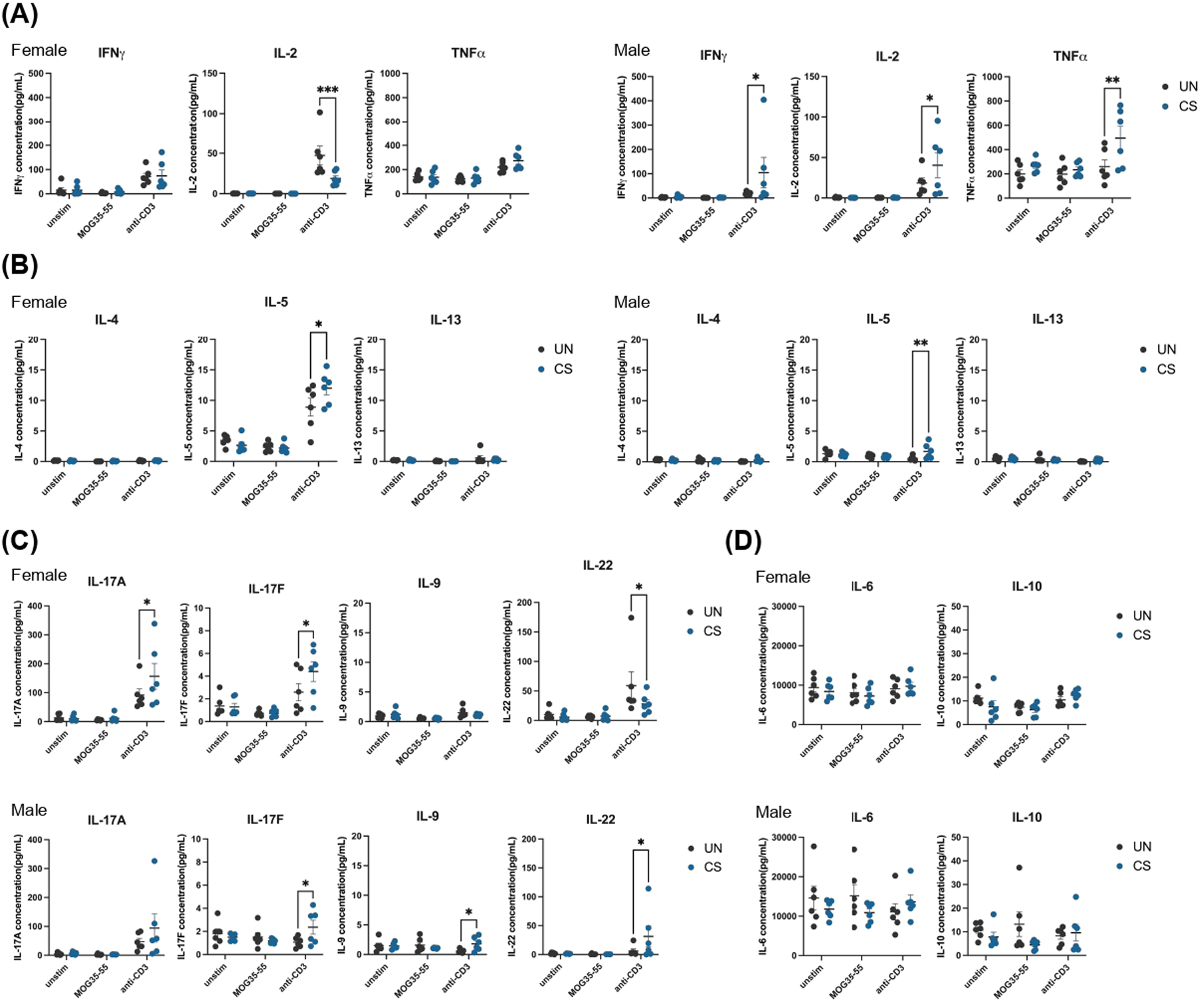
Sex-specific T lymphocyte responses to chronic stress. Eight-week-old C57BL/6 mice were either left unstressed (UN) or subjected to 8-wk UCMS (CS). Whole brains were dissected, enzymatically digested and cells isolated by percoll gradient followed by in vitro restimulation with MOG_35-55_ peptide or anti-CD3 antibody for 72 hours. Supernatant was collected at the endpoint and cytokine concentration measured. Raw results were normalized with CD45+ cell counts. **(A)** IFNγ, IL-2 and TNFα concentrations. **(B)** IL-4, IL-5 and IL-13 concentrations. **(C)** IL-17A, IL-17F, IL-9 and IL-22 concentrations **(D)** IL-6 and IL-10 concentrations. Data are shown as mean ± SEM. Two-way RM ANOVA with Tukey’s multiple comparisons test was used. * p < 0.05; ** p < 0.01. Each replicate corresponds to an individual mouse.

Altogether, these results reinforce the increase of male-specific type 1 signature in brain lymphocytes in chronic stress also observed in the scRNAseq data. Type 17 cytokines were found elevated in chronic stress in both sexes, with a stronger IL-17A and IL-17F signature in females and a more IL-9, IL-22 oriented response in males. Based on the scRNAseq data, these type 17 signatures induced in chronic stress conditions, could be attributed to rγδT17 cells, further emphasizing their sex-dimorphic role in disease context.

## 4. Discussion

Women and men differ significantly in both the prevalence and clinical presentation of depression (Salk et al. 2017; Eid et al. 2019). At the molecular level, this disparity is supported by sex-specific genetic architectures associated with depression (Thomas et al. 2025) and by the differences in the hypothalamus-pituitary-adrenal axis function in stress (Heck and Handa 2019; Panagiotakopoulos and Neigh 2014). Immune responses are likewise sexually dimorphic, raising the possibility that sex-specific immunity contributes to the observed bias in depression and stress-related pathology (Klein and Flanagan 2016; Tukiainen et al. 2017; Sharma et al. 2025; Bekhbat and Neigh 2018; Lenz and McCarthy 2015). Defining these immune mechanisms may reveal molecular targets for sex-tailored treatments for depression, a disorder that already exhibits sex differences in symptom profile and severity.

Unfortunately, males have been predominantly used in preclinical research, leading to insufficient consideration of sex differences in studying immune responses in stress biology. Part of this bias comes from model limitations: for example, in the social defeat model, which is widely used to study social aggression, can only be applied to male rodents. However the development of model variation now enables female analysis in social defeat models (Takahashi et al. 2017), and has helped reveal stress mechanisms unique to females (Dion-Albert et al. 2022). The current study uses the unpredictable chronic mild stress model which effectively induces depressive-like behaviors in both females and males. We analyzed stress-and sex-dependent alterations in behaviors, hippocampal myelin integrity, neuronal structure, and the overall immune cell landscape of the brain. Sex bias was evident across all aspects of our analysis. Notably, pronounced sex differences emerged in the composition and phenotype of brain-resident lymphocytes: males exhibited a higher frequency of NKT and αβT cells, whereas females displayed an increased proportion of γδT cells. Overall, female lymphocytes have a significantly higher enrichment of type 2 and 17 responses as well as higher expression of inflammasome-related genes, as compared to males (Figure 5B, C and G). The sex disparity is further exacerbated when mice were exposed to UCMS (Figure 5B and C). Importantly, although chronic stress increased IL-17F expression in both sexes, IL-22 expression diverged: rising in males but declining in females, reflecting distinct polarizations within type 17 responses (Figure 6C). Together, these findings demonstrate that sex-specific immune biases manifest at both cellular and molecular levels and are amplified during stress. Future functional and anatomical studies will be essential to determine how these immune differences contribute to sex-specific depressive pathology, such as the hippocampal changes observed in this study.

While chronic stress affects the hippocampus in both sexes, the nature and extent of these alterations are sex-dependent and may underlie divergent behavioral outcomes, particularly for anxiety and social behaviors. The structural consequences of chronic stress on the hippocampus are multifaceted, encompassing histological, molecular and structural changes. These include region-specific gray matter atrophy, as well as reductions in neuronal number and synaptic density (Roddy et al. 2019; Sheline 2011; Nunes et al. 2022; Mechawar and Savitz 2016). Stress exposure is associated with decreased expression of key synaptic vesicle proteins, including PSD95 and SYN, which correlate with diminished dendritic spines and branching (Fuchs et al. 2006; Shen et al. 2021; 2019). Together, these neuronal changes also collectively impair hippocampal functional connectivity, with emerging evidence indicating sex-dependent differences in both the magnitude and pattern of these disruptions (Negrón-Oyarzo et al. 2016; Narayanan and Chattarji 2010; Jefferson et al. 2020).

In this study, we recapitulated key alterations in hippocampal organization following UCMS exposure. Female mice exhibited marked CA1 atrophy and reduced neuronal layer thickness in the CA2, and DG regions. Consistently, PSD95 levels were decreased in hippocampus of UCMS-exposed females but remained unchanged in males, while SYN expression was unaffected in both sexes. Although our model does not capture all aspects of hippocampal organization disruptions reported in literature, our findings collectively indicate that UCMS induces widespread structural remodeling of the hippocampus, with greater vulnerability in females.

The neuronal alterations observed in our study are not associated with myelin dysregulation, unlike those reported in studies of human depression (Tham et al. 2011; Sacchet and Gotlib 2017; Baranger et al. 2021; Shim et al. 2023). As prior work has documented corpus callosum structural abnormalities (Ran et al. 2020; Cole et al. 2012), and cortical demyelination in both humans (Tham et al. 2011) and rats (Yang et al. 2015), it remains possible that myelin integrity in regions not analyzed in this study can be affected. In fact, both IL-17 and IFNγ have been reported to play a significant role in regulating oligodendrocytes and precursors (Liu et al. 2021; Wang et al. 2017; Horiuchi et al. 2006; Kirby et al. 2019; Molina-Gonzalez et al. 2022), as well as impacting neuron health, neurogenesis and axonal regeneration (Lee et al. 2025; Brigas et al. 2021; Cristiano et al. 2019; Liu et al. 2014; Wang et al. 2023). As such, the chronic-stress-dependent induction of IL-17 and IFNγ can play a direct role in the sex- and region-dependent neuronal and myelin dysregulation.

Our findings support that IL-17 and IFNγ in the brain originate from tissue-resident lymphocytes, γδT versus αβT and NKT cells, respectively, a finding consistent with previous report of meningeal lymphocytes (Alves de Lima et al. 2020). The presence of distinct lymphocyte lineages producing diverse cytokines within the brain is intriguing and may reflect the nature of antigens such as lipid, oligosaccharides, protein, and microbial components, encountered either locally in the brain or in distal tissues such as the gut and adipose tissues. We did not detect any expression of Vγ7, a selective γ chain used by intestinal epithelial lymphocytes. Further, while we found enriched MR1 expression, a receptor for microbial product, the MR1^+^ αβT cells do not have a restricted α or β chain usage as described for MAIT cells at the mucosal sites. While our analysis does not support an intestinal origin of brain-resident lymphocytes, future studies using direct approaches are needed to rule out this possibility.

Tissue resident memory T cells are increasingly recognized as key players of infection, cancer and autoimmune conditions (Sasson et al. 2020; Xie et al. 2025; Christo et al. 2024). Our data support that UCMS promoted cytokine responses in different lymphocyte lineages. Whether the production of IL-17 and IFNγ by tissue resident lymphocytes in response to UCMS plays a protective or detrimental role in the pathogenesis of depression remains unclear. Clinically, blockade of upstream signals such as IL-12 and IL-23 has significantly reduced depression symptoms in an array of inflammatory diseases (Wittenberg et al. 2020). IFNγ promotes the activation of immunoproteasome and remodels immunopeptidome (Newey et al. 2023) and as such, may diversify the Tc1 responses in the UCMS brain leading to increased risks of inflammation. Conversely, endogenous IFNγ and IL-17 have been implicated in maintaining normal social interaction behaviors (Filiano et al. 2016) and anxiety regulation (Alves de Lima et al. 2020) suggesting that their upregulation under chronic stress may represent an adaptive mechanism to counter depressive-symptoms. Notably, UCMS induced distinct type 17 signatures in males, characterized by increased IL-9 and IL-22 expression. The functional roles of these and other cytokines in the neuroimmune response to chronic stress and depression warrant further investigation.

How the UCMS-dependent sex-specific expansion of type 1 and type 17 responses is regulated remains an important question. Our data suggest that female and male lymphocytes differentially suppressed ZFP36 and P4HA1. ZFP36/TTP is a translational suppressor that recruits mRNA to stress granules (Fu and Blackshear 2017), a membraneless organelle induced by cellular stress (Wolozin and Ivanov 2019; Cui et al. 2024). We found that chronically stressed female lymphocytes reduced the expression of TTP but not any of the other family members ZFP36L1 and ZFP36L2 which have preferential binding capacity to different types of mRNA (Makita et al. 2021). TTP preferentially binds to mRNA of various inflammatory cytokines (Fu and Blackshear 2017; Makita et al. 2021). Previous work show that TTP in T cells restrains IL-17 production in aged mice (Peng et al. 2020). Thus, we hypothesize that the female brain resident lymphocytes have expanded IL-17 response due to the loss of TTP expression. On the other hand, male lymphocytes suppressed P4HA1 expression in UCMS. P4HA1 is a hypoxia-induced protein that accumulates and impairs mitochondrial fitness by disrupting α-ketoglutarate metabolism in the tricarboxylic acid cycle (Ma et al. 2025). α-ketoglutarate is well-established for regulating the metabolism and function of Tregs, CD8 T cells, and Th17 cells (Minogue et al. 2023; Klysz et al. 2015; Matias et al. 2021; Wagner et al. 2021). As such, we reason that the loss of P4HA1 expression in male UCMS brain lymphocytes contributes to the exacerbation of IFNγ, TNFα, and IL-2 expression. This hypothesis is partly supported by our *ex vivo* cytokine analysis shown in Figure 6.

In sum, this study has identified chronic-stress- and sex-driven molecular and cellular changes in brain resident lymphocytes, hippocampal synaptic organization, and depressive-like behaviors, providing a framework from which disease pathogenesis of depression as well as other conditions primed by chronic stress can be functionally tested.

## Supporting information

Raw Western Blot images of Figure 2 and Supplementary Figure 2

## 6. Author Contributions

SP, KC, and CW conceived the project. SP, KC and CW wrote the manuscript with help from YP, SW and AT. SP performed and analyzed immune cytokine and western blot experiments and coordinated all hippocampal pathology analysis. KC performed the scRNASeq experiments and performed all the relevant data analysis. YP performed the IHC experiments and analysis. SW acquired and analyzed the SRS microscopy data under the supervision of CQ. KJL established the UCMS model. KJL and MN performed behavioral tests with help from DU. DU also performed the corticosterone experiments and helped with cell isolation. AT performed the UCMS experiments, extracted tissue and cell for downstream analysis and edited the manuscript. CW supervised the study.

## 8. Supplementary Material

Supplementary material include supplementary figures 1 to 5 and supplementary table 1 to 3. Raw images of the western blots presented in Figure 2 and Supplementary Figure 2 are also available.

## 9. Resource Availability Statement

This study does not report original code. Single-cell RNA sequencing data and any additional information required to reanalyze the data reported in this paper is available from the lead contact upon request. Lead contact: Please direct requests for further information and resources to Chao Wang. No new reagents were generated.

---

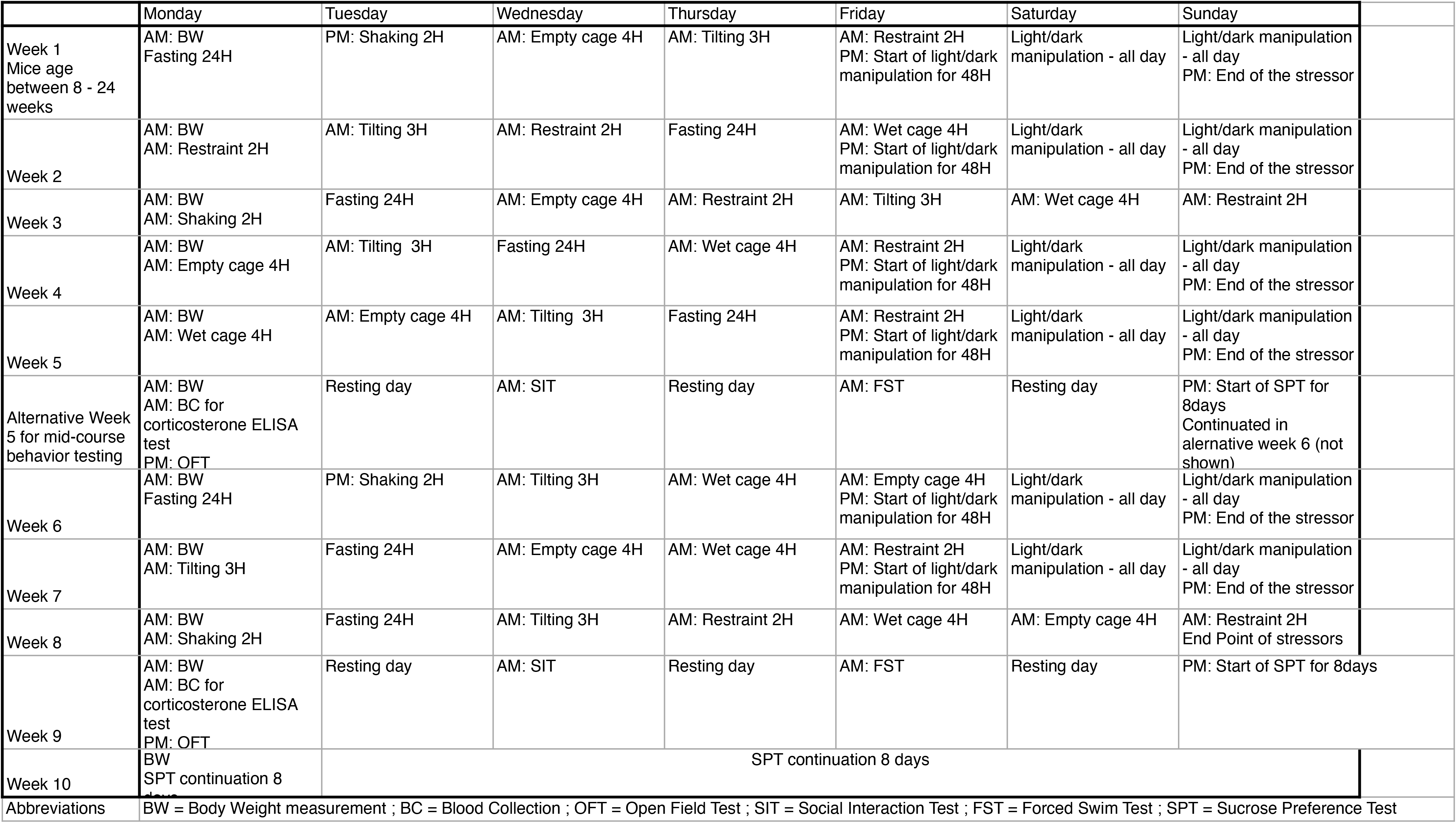

