## Supplementary Figures 1-5 for "Chronic stress promotes sex-divergent Type 1, 2 and 17 responses in brain resident lymphocytes"

### *Supplementary Material*

#### 1. Supplementary Figures

(A)

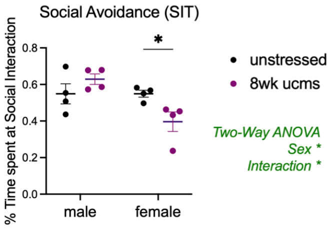


**Supplementary Figure 1.** Same experimental set up as in Figure 1**. (A)** Social interaction test was performed for age-matched unstressed and 8-week UCMS mice. Each dot is a mouse. Data was analyzed by two-way ANOVA with Tukey's multiple comparisons test. * $p < 0.05$.

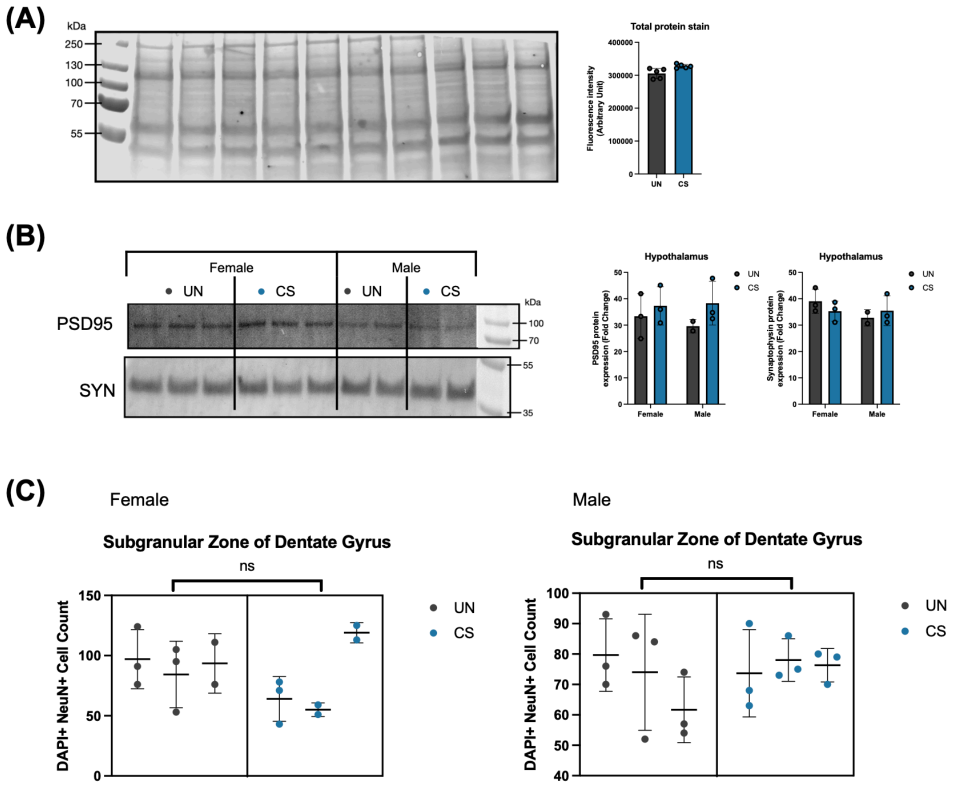


**Supplementary Figure 2.** Eight-week-old C57BL/6 mice were either left unstressed (UN) or subjected to 8-wk UCMS (CS). **(A)** Exemple of total protein quantification used to normalize the results of protein expression in Western Blot. **(B)** PSD95 and SYN expression levels measured by Western Blot in the hypothalamus. Data are shown as mean ± SD. Two-way ANOVA with Fisher's LSD multiple comparisons test was used. **(C)** Quantification of NeuN+ DAPI+ double positive cells in the Subgranular Zone of Dentate Gyrus. Data is presented as mean ± SD and analyzed using nested t-test.

**(A)**

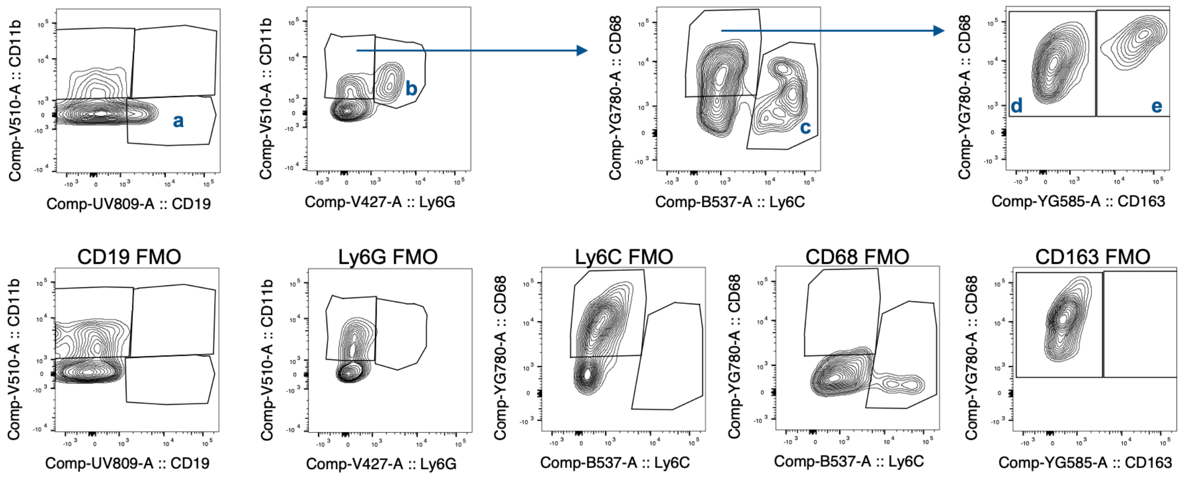


**(B)**

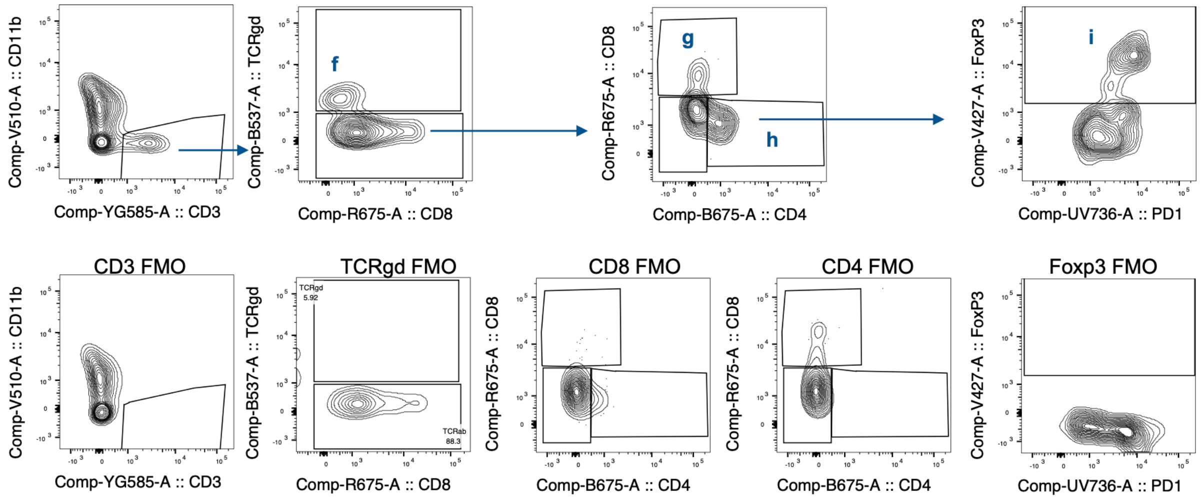


**Supplementary Figure 3.** Flow cytometry gating strategy for Figure 3. Cells were stained for either myeloid panel **(A)** or T cell panel **(B)** and analyzed by BD Symphony A5 gated on FSC vs SSC, single cells, and $CD45^{high}$. FMO: fluorescence minus one control. Each identified population is as follows: a. B cells ($CD45^+CD19^+$); b. Neutrophils ($CD45^+CD11b^+Ly6G^+$); c. $Ly6C^+$ myeloid cells ($CD45^+$ $CD11b^+$ $Ly6C^+$); d. Macrophage ($CD45^+CD11b^+CD68^+CD163^-$); e. Border Associated Macrophage (BAM, $CD45^+$ $CD11b^+CD68^+CD163^+$); f. gamma delta T cells ($CD45^+CD11b^-CD3^+TCRgd^+$); g. CD8 T cells ($CD45^+$ $CD3^+TCRgd^-CD8^+$); h. CD4 T cells ($CD45^+CD3^+TCRgd^-CD4^+$); i. $CD4^+$ Tregs ($CD45^+CD3^+TCRgd^-CD4^+Foxp3^+$).

**(A)**

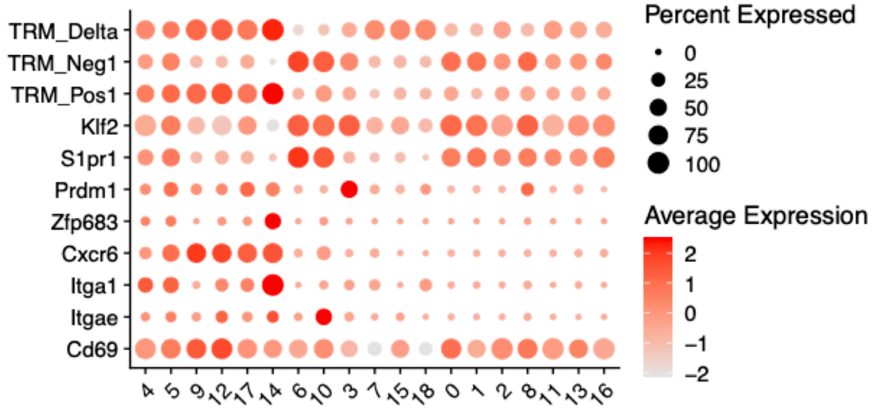


**(B)**

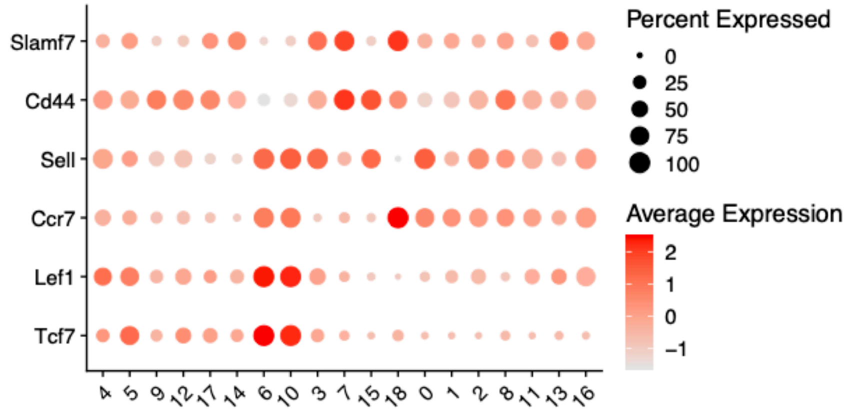


**Supplementary Figure 4. Expression of tissue-resident and activation-associated markers in lymphocyte clusters. (A)** Dot plot showing expression of canonical tissue-resident lymphocyte markers. **(B)** Dot plot showing expression of naïve-associated and activation-related markers across clusters.

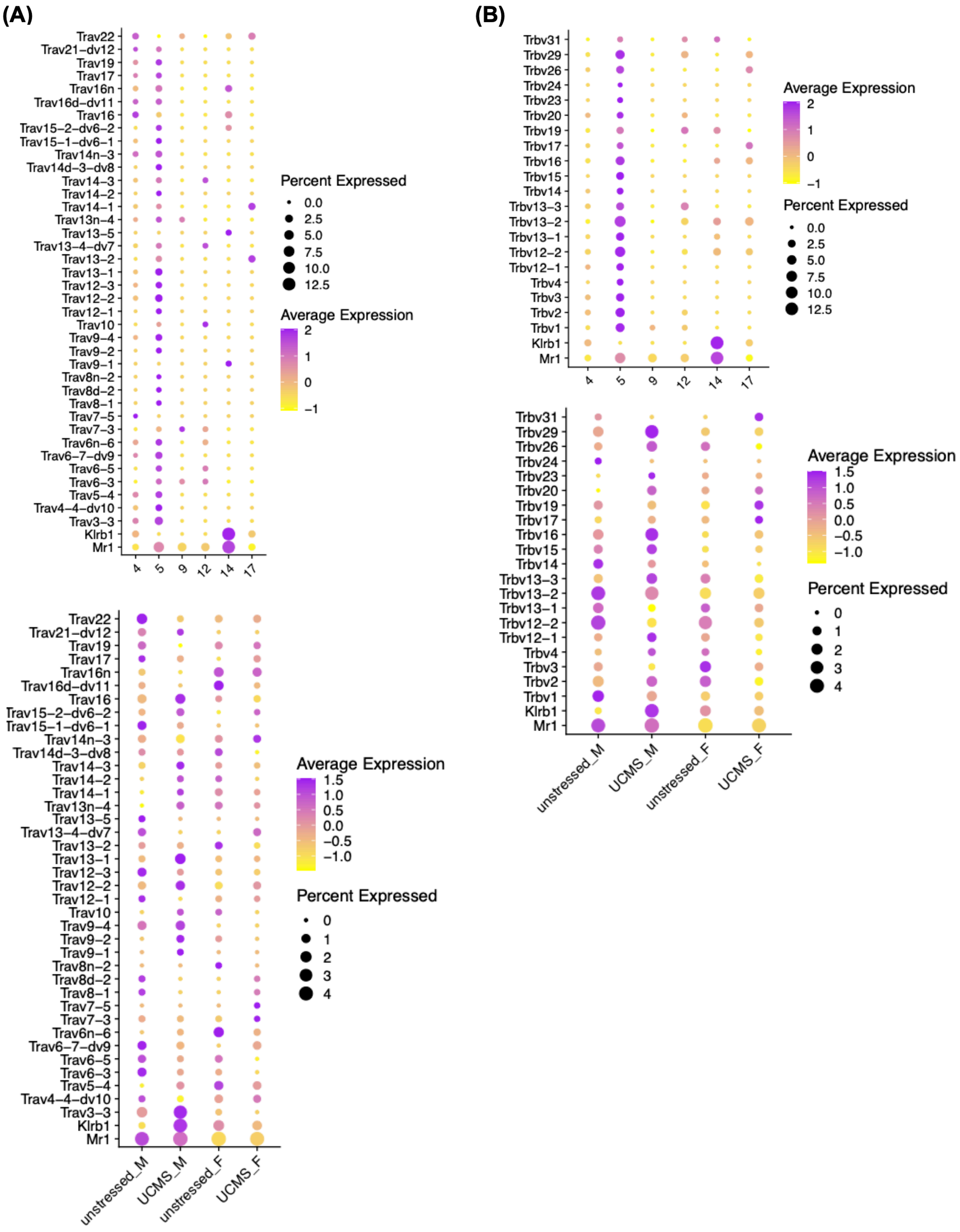


**Supplementary Figure 5. Expression of T-cell receptor variable region genes in MAIT cells.** Dot plot showing gene expression of Mr1, Klrb1, and **(A)** Trav family or **(B)** Trbv family across tissue-resident lymphocyte clusters and conditions.

#### 2. Supplementary Tables

**Supplementary Table 1. Marker genes for lymphocyte clusters.** All differentially expressed genes were identified using Seurat's FindAllMarkers function across lymphocyte clusters at resolution 0.5.

**Supplementary Table 2. Differentially expressed genes between female and male in tissue-resident lymphocytes.** Differential expression analysis was performed on tissue-resident lymphocytes using Seurat's FindMarkers function, comparing females (F) and males (M).

**Supplementary Table 3. Differentially expressed genes in tissue-resident lymphocytes by stress condition.** Differential expression analysis was performed on tissue-resident lymphocytes using Seurat's FindMarkers function, comparing UCMS and unstressed conditions separately in males (UCMS_M vs unstressed_M) and females (UCMS_F vs unstressed_F).
