## Supplementary material for "Chronic stress promotes sex-divergent Type 1, 2 and 17 responses in brain resident lymphocytes": Raw Western Blot images of Figure 2 and Supplementary Figure 2

**Figure 2. UCMS alters hippocampal neuron organization but not myelin integrity.** Eight-week-old C57BL/6 mice were either left unstressed (UN) or subjected to 8-wk UCMS (CS). Hippocampi were processed either for protein extraction or for immunohistochemistry. **(A)** Western Blot showing PSD95 and Synpatophysin (SYN) with their protein expression normalized using Total Protein Quantification. Data are shown as mean ± SD. Two-way ANOVA with Fisher's LSD multiple comparisons test was used. Each replicate corresponds to an individual mouse. **(E)** Western Blot showing Myelin-Basic Protein (MBP) expression in female mice and its protein expression normalized using Total Protein Quantification.

(A) PSD95 staining - full membrane – post LUT-inversion

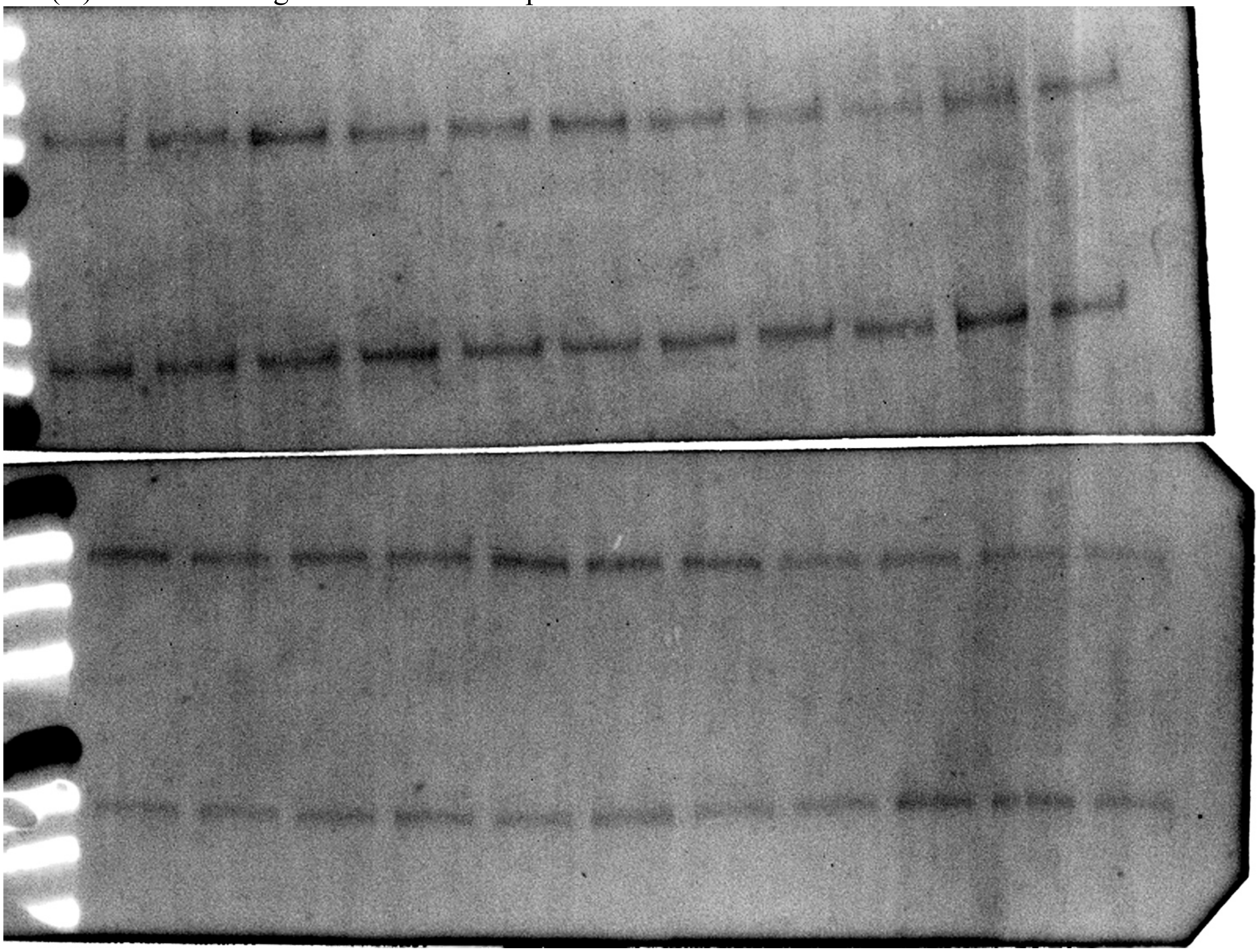

(A) Synaptophysin staining - full membrane post LUT-inversion

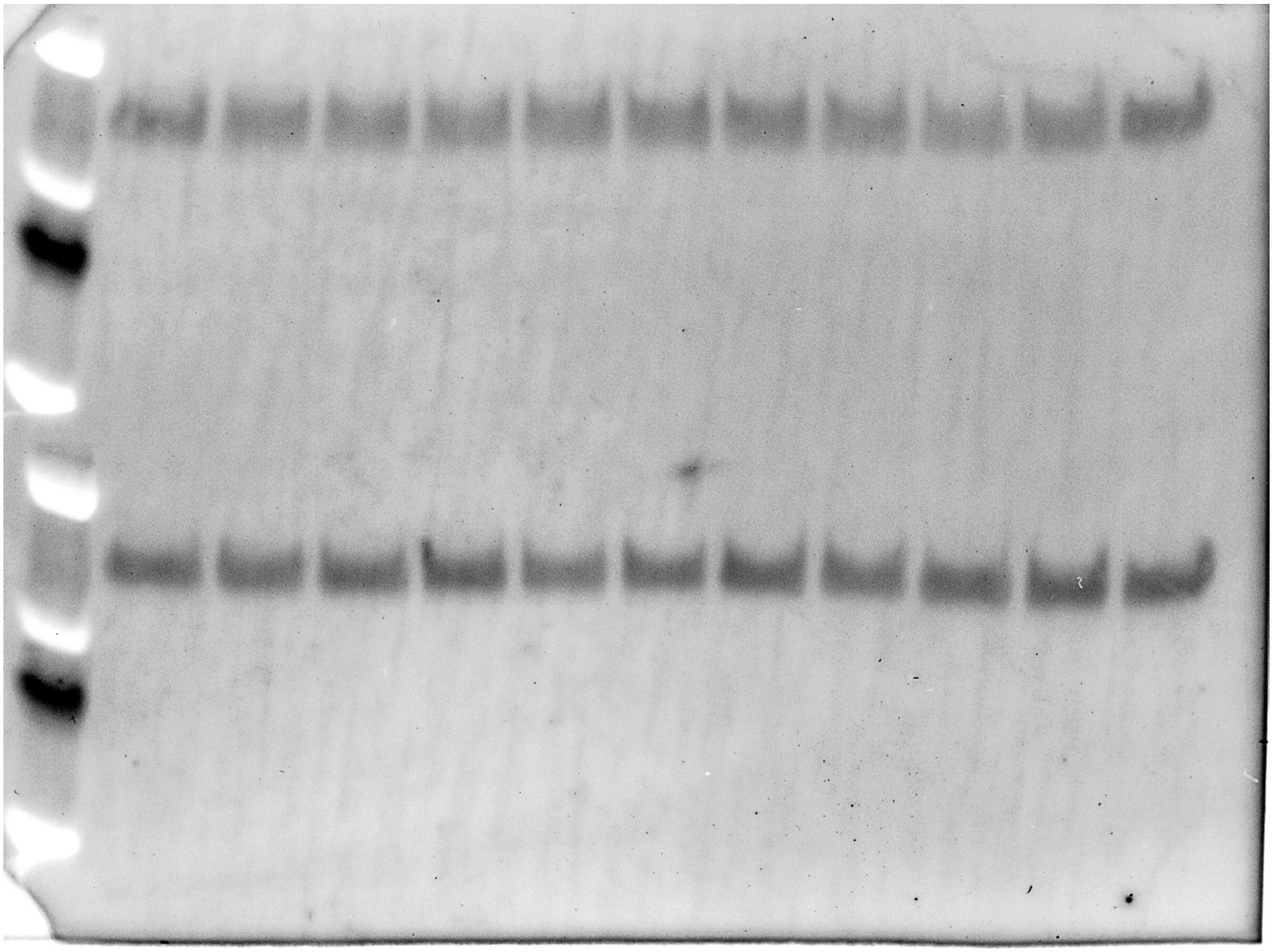

**(E)** MBP staining - full membrane post LUT-inversion

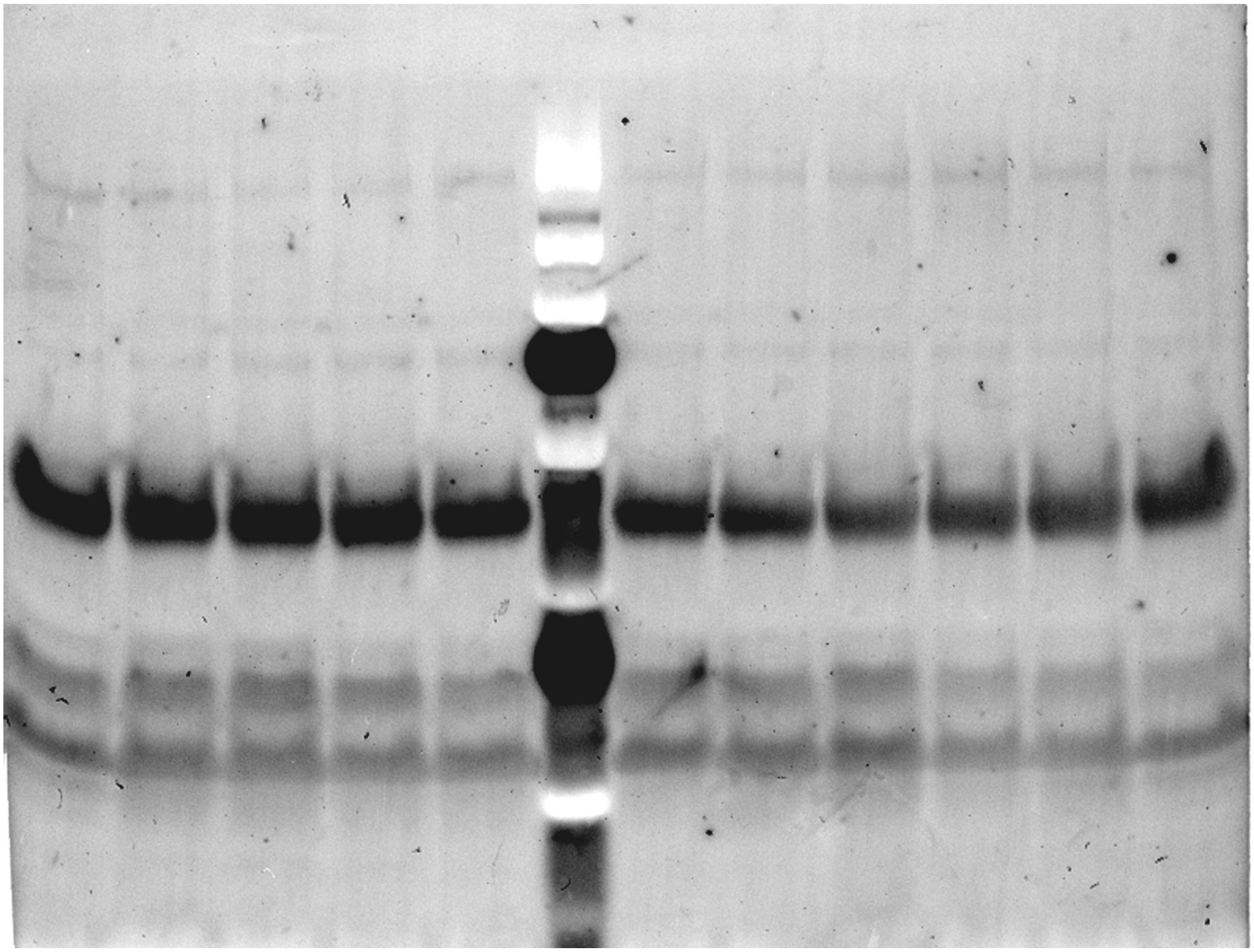

**Supplementary Figure 2.** Eight-week-old C57BL/6 mice were either left unstressed (UN) or subjected to 8-wk UCMS (CS). **(A)** Example of total protein quantification used to normalize the results of protein expression in Western Blot.

**(A)** Total protein quantification – full image of 2 membranes

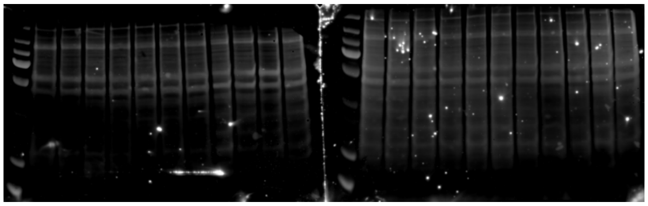
